# The appearance of *sugI* mixed loci in three individuals during treatment for MDR-TB, supports the involvement of *sugI* in *Mycobacterium tuberculosis* d-cycloserine resistance *in vivo*

**DOI:** 10.1101/2023.05.30.542839

**Authors:** R.M Anthony, M. Molemans, O. Akkerman, M.G.G. Sturkenboom, A. Mulder, R. de Zwaan, D. van Soolingen, J. Alffenaar, F.C.M. van Leth, S. Ghimire, N. Yatskevich, A. Skrahina, N. Ciobanu, N. Turcan, V. Crudu

## Abstract

To study the adaptation of multi-drug resistant *Mycobacterium tuberculosis* (MDR-TB) during treatment patients diagnosed with MDR-TB were recruited into an observational study. Clinical data and *M. tuberculosis* DNA at diagnosis and between seven days and two months of MDR-TB treatment were collected. The drugs prescribed were recorded. Interpretable WGS data from 118 isolates from 54 participants was obtained (11 in Belarus and 43 in Moldova) and screened for the presence of unfixed single nucleotide polymorphisms (mixed SNPs / loci).

This study was performed shortly after the publication of the 2019 WHO consolidated guidelines on drug-resistant tuberculosis treatment. Existing drug supplies and procurement in one country after the switch to the all oral MDR-TB regimen in addition to patient factors, influenced the selection of and exposure to drugs.

Confidently mixed SNPs were identified in samples from multiple participants in only five genes (*gyrA, pncA*, Rv1129c, Rv1148c, and *sugI*). All other genes with confidently mixed SNPs were identified in isolates from only a single individual. A significant proportion of the participants (52 of 54 participants) received d-cycloserine as part of their initial treatment, most participants who initially received d-cycloserine did not receive bedaquiline in their initial regimen (all at one site). Three different mixed SNPs were identified in *sugI* gene from a follow up isolate from three participants (P7A, P7T, and Q6stop). Mutations in *sugI* have previously been reported in spontaneous *in vitro* d-cycloserine resistant mutants. Alterations in the *sugI* gene may indicate a sub optimal d-cycloserine containing regimen and potentially be of clinical significance with respect to adaptation to d-cycloserine. Monitoring the accumulation of low frequency escape mutants may help identify regimens insufficiently powerful to block the accumulation of antimicrobial resistance mutants and identify drug(s) at risk of resistance selection.

## Introduction

Routine genotyping of *M. tuberculosis* isolates by whole genome sequencing (WGS) generally relies on mapping to a reference genome and only calling SNPs when they represented the majority of the mapped reads (typically >70-80%). Lower frequency SNPs are more difficult to identify and interpret and are only routinely called when specifically screened. Loci screened for low frequency mutations are in most pipelines confined to positions with a known (strong) association with drug resistance [Anthony 2023] or for SNPs characteristic for major lineages that if mixed suggest the presence of more than one mycobacterial genotype [Coll 2014].

Nonetheless the accumulation of non-fixed mixed SNPs during treatment has been observed and is associated with the *in vivo* emergence of resistance in an individual patient [Castro 2021, Nimmo 2020]. Not all antibiotic escape mutations are successful and multiple resistance mutations associated with a single drug have been reported [Trauner 2017]. This accumulation of mutations may be an indicator for a “fragile” regimen [Fox 2022] where the concentration / number of active of drugs is insufficient to prevent the accumulation of resistance mutations. Furthermore the particular spectrum of mutations seen might indicate the key drug(s) where selective pressure is being most strongly applied to the mycobacteria by the chosen regimen. In an attempt to explore these phenomena we performed WGS on serial isolates form participants treated for drug resistant tuberculosis, screened the sequences produced for mixed SNPs and recorded the drugs prescribed.

## Methods

### Patient recruitment

The study was conducted across multiple treatment centers in Europe, including the Institute of Phthisiopneumology in Chisinau, Moldova; TB-Net in Minsk, Belarus; and three centers in the Netherlands. The Dutch centers comprised the National Tuberculosis Reference Laboratory at the Centre for Infectious Disease Control, the Amsterdam Institute for Global Health and Development (AIGHD), and the University Medical Center Groningen. The coordinating center for the study was the National Institute for Public Health and the Environment (RIVM) in the Netherlands, which served as the primary point of contact. The study protocol was submitted to the Medical Ethical Review Board in Groningen (METc Groningen), and was deemed exempt from the Medical Research Involving Human Subjects Act by METC (2019/354). Between July 2019 and January 2022 patients diagnosed with MDR-TB, on the basis of a rapid screen for rifampicin resistance or clinical suspicion, were informed about the study those giving their informed consent were recruited.

### Clinical data

Sociodemographic characteristics and the regimen received by the participants were collected in Open Data Kit using tablets provided to the clinical sites, with electronic questionnaires at three time points: day 0, day 14 (window 7-11 days), one month (window 28-35 days) and at 2 months (window 58-62 days) (Supplementary table 1). Data validation with queries was performed during the study. Data analysis was conducted in Stata 16.1.

### Screening for non-fixed / mixed SNPs

DNA was extracted from cultured mycobacteria by heat lysis and purified on QIAamp DNA Mini Kit spin columns (Qiagen Düsseldorf, Germany) in the isolating laboratories. DNA was then shipped to the Netherlands and subjected to whole genome sequencing (WGS) using Illumina (CA, USA) (HiSeqTM 2500, NextSeqTM 500 or NextSeqTM 550 sequencer) and mapped to the reference genome (GenBank accession: AL123456.3) [Jajou 2019]. The FASTQ files (NCBI BioProject 825716 https://www.ncbi.nlm.nih.gov/sra?linkname=bioproject_sra_all&from_uid=825716) were then analysed by LoFreq [Wilm 2012] and loci annotated according to the reference genome (https://github.com/RIVM-bioinformatics/Myco_lofreq). The resulting CSV files were then filtered with an R-script to remove calls with a LoFreq quality score <150 or within 6 bp of another single nucleotide polymorphism (SNP). This resulted in a list of 200 to 350 loci with mixed SNP calls (allele frequency 0.05% to 0.98%) for each of the samples analyzed.

The majority of the 200-350 mixed SNPs even after filtering when visually inspected (SAMTools “pileup” [Li 2009]) were found to be present in complex poorly mapped regions (e.g. PE/PPE) or likely multi copy regions one or more of which contained fixed SNPs and considered unlikely to represent real mixed SNPs. It was also observed that the majority of these mixed SNPs were called repeatedly in unrelated samples, further suggesting they were sequencing artifacts. In order to streamline the analysis these regions and a supplementary list of SNPs (176 positions, supplementary table 2) that were called in multiple unrelated samples and considered unreliable based on visual inspection of the SAMTools pileups were used to filter the non-fixed SNP lists obtained from the individual isolates. The SAMTools pileups from the remaining shorter lists of non-fixed SNPs from each isolate were then visually inspected and only confident calls retained. This list (supplementary table 4) was then screened for genes containing mutations in multiple patients and mutations in any genes reported to be associated with resistance to antimycobacterial drugs (supplementary table 5). Finally, the Rv0678 gene was screened for Indels (Supplementary table 6).

## Results

### Patient recruitment

As a result of political instability it was unfortunately not possible to ship all samples collected for analysis (Figure 1). Twenty five DNA extracts from 11 eligible participants recruited in Belarus were collected and shipped for analysis (all treated as MDR although one was initially RIF sensitive based on WGS and a locally performed LineProbe MDR-TB plus [Hain diagnostics, Germany] but this patient had complex treatment history so a clinical decision was made to treat this patient with an MDR-TB regimen). One hundred DNA samples were collected and shipped from Moldova 93 of these samples from 43 participants were analyzed for mixed loci. Seven samples were excluded: two samples from 2 participants as they were not diagnosed/treated as MDR-TB, four samples had a different genotype to the initial isolate, and one sample had a mixed genotype based on the WGS analysis (Figure 1).

**Figure 1:**
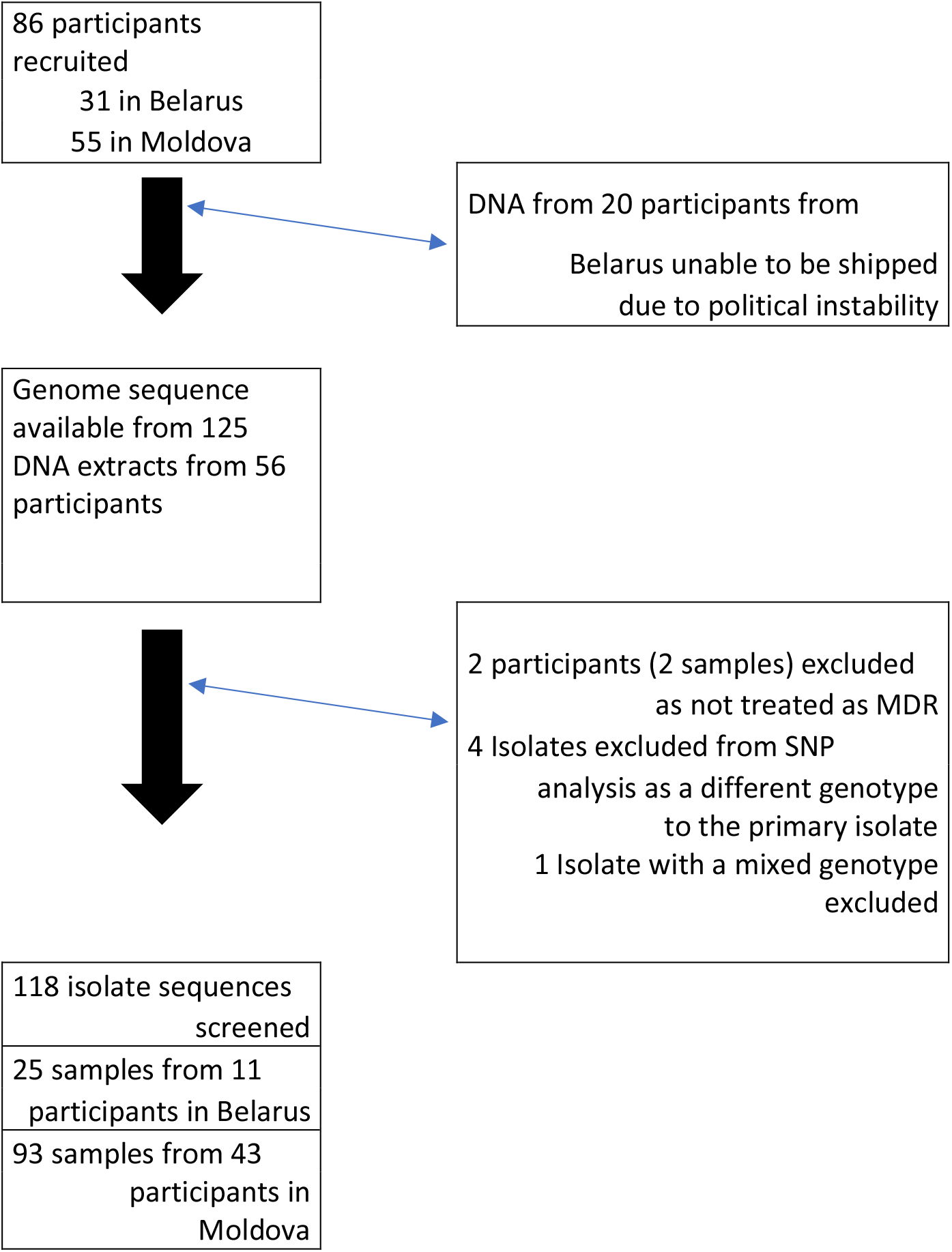
Participant and sample flowchart of samples shipped and analysed by WGS for unfixed SNPs (a list of all sequence files and their origin is available in supplementary table 3).

### MDR-TB treatment prescribed

The drugs prescribed were recorded (supplementary table 1) and drugs provided at the time points the samples were collected summarized in Table 1.

**Table 1.** Summary of the drugs prescribed at the time points when the cultures (DNA extracts) were collected during the study.

| Site | Follow up (number of participants on treatment) | Number (%) of participants with drug in Initial regimen |  |  |  |  |  |  |  |  |  |
| --- | --- | --- | --- | --- | --- | --- | --- | --- | --- | --- | --- |
|  |  | BDQ | CYS | LFX | MFX | LZD | CFZ | DLM | IMI | AMO | PAS |
| Belarus | Diagnosis (11) | 11(100) | 11(100) | 9(82) | 0(0) | 11(100) | 11(100) | 1(9) | 1(9) | 1(9) | 0(0) |
|  | Seven days (11) | 11(100) | 11(100) | 9(82) | 0(0) | 11(100) | 11(100) | 1(9) | 1(9) | 1(9) | 0(0) |
|  | One month (11) | 11(100) | 11(100) | 9(82) | 0(0) | 11(100) | 11(100) | 1(9) | 1(9) | 1(9) | 0(0) |
|  | Two Months (11) | 11(100) | 11(100) | 9(82) | 0(0) | 11(100) | 11(100) | 1(9) | 1(9) | 1(9) | 0(0) |
| Moldova | Diagnosis (43) | 5(12) | 41(95) | 34(79) | 3(7) | 41(95) | 2(5) | 0(0) | 13(30) | 13(30) | 14(33) |
|  | Seven days (42) | 6(14) | 40(95) | 33(79) | 3(7) | 41(98) | 2(5) | 0(0) | 15(36) | 15(36) | 18(43) |
|  | One month (42) | 15(36) | 40(95) | 33(79) | 3(7) | 41(98) | 3(7) | 0(0) | 16(38) | 17(40) | 15(36) |
|  | Two Months (41) | 19(46) | 39(95) | 30(73) | 2(5) | 39(95) | 3(7) | 0(0) | 15(37) | 16(39) | 16(39) |

BDQ = bedaquiline, CYS = d-cycloserine, LFX = levofloxacin, MFX = moxifloxacin, LZD = linezolid, CFZ = clofazimine, DLM = delamanid, IMI = imipenem-cilastatin, AMO = amoxicillin-clavulanic acid PAS = *p*-aminosalicylic acid, PZA = pyrazinamide, Ethp = ethionamide-prothionamide, CAP = capreomycin

### Screening for Mixed SNPs

The data generated is available as NCBI bioproject PRJNA825716 and described in supplementary table 3). Initial screening revealed, one sample (MD-17-12) contained a mixed genotype, based on the presence of large number of mixed SNPs (>500) with a similar frequency [de Neeling 2022], for four isolates (MD27-14, MD46-13, MD42-12, MD52-13) the genotype of the isolated strain was different to the initial isolate (>20 SNPs), two patients did not receive treatment for MDR-TB (MD23 and MD 29), these samples were excluded (Figure 1). Sequencing data consistent with a single strain was analyzed from the 118 isolates from 54 participants. Between 11 and zero confident unfixed / mixed (>5%) SNPs were identified in the Ilumina (CA, USA) data from the isolates examined with an average of 4 mixed SNPs identified from each patient (Figure 1, Supplementary table 4).

The WGS data was analyzed blinded to the prescribing data. Mixed SNPs present in the same gene from multiple participants were identified in only five genes; *gyrA, pncA*, Rv1129c, Rv1148c, and *sugI* in two, two, three, two and three participants respectively (Table 2). All other genes with mixed SNPs were present only in a single patient (Supplementary table 4). Non-fixed indels were not screened in the whole genome although the Rv0678 gene was examined in more detail as it is known to be associated with bedaquiline resistance [Villellas 2017, Nguyen T.V.A 2018], variability was detected in two participants. Multiple mixed Rv0678 insertions were detected in one patient (779117 ins T, and 779182 ins G in patient MD-52) a follow up isolate had the 779117 T insertion fixed (>95% of the reads) (Supplementary table 6). The Rv0678 gene also contained a mixed SNP in a second patient (778994 G>T Rv0678 S2I in patient MD-16, 10% of the reads in the first isolate and 34% of the reads in the second isolate, supplementary table 5).

**Table 2:** Mixed loci in genes that contained confidently mixed SNPs in more than one patient.

| County-Participant-sample | Genome position (H37Rv) | Mutant proportion | Read depth | Original codon | Mutant codon | AA change | Gene |  |
| --- | --- | --- | --- | --- | --- | --- | --- | --- |
| MD-011-13 | 8416 | 0.22 | 315 | GTC | GCC | V327A | Rv0006 | *gyrA* |
| MD-047-11 | 8208 | 0.82 | 373 | GAG | CAG | E303Q | Rv0006 | *gyrA* |
| MD-047-12 | 8208 | 0.99 | 304 | GAG | CAG | E303Q | Rv0006 | *gyrA* |
| MD-016-11 | 2288920 | 0.76 | 133 | GGA | CGA | G108R | Rv2043c | *pncA* |
| MD-016-11 | 2289081 | 0.05 | 119 | CCG | CTG | P57L | Rv2043c | *pncA* |
| MD-016-12 | 2288920 | 0.56 | 419 | GGA | CGA | G108R | Rv2043c | *pncA* |
| MD-016-12 | 2289081 | 0.47 | 412 | CCG | CTG | P57L | Rv2043c | *pncA* |
| MD-040-11 | 2289213 | 0.81 | 346 | CAG | CGG | Q10R | Rv2043c | *pncA* |
| MD-040-12 | 2289213 | 0.90 | 274 | CAG | CGG | Q10R | Rv2043c | *pncA* |
| MD-040-13 | 2289213 | 0.77 | 221 | CAG | CGG | Q10R | Rv2043c | *pncA* |
| MD-040-14 | 2289213 | 0.86 | 236 | CAG | CGG | Q10R | Rv2043c | *pncA* |
| MD-040-11 | 1254318 | 0.82 | 385 | GCG | ACG | A73T | Rv1129c | *prpR* |
| MD-040-12 | 1254318 | 0.91 | 380 | GCG | ACG | A73T | Rv1129c | *prpR* |
| MD-040-13 | 1254318 | 0.80 | 264 | GCG | ACG | A73T | Rv1129c | *prpR* |
| MD-040-14 | 1254318 | 0.92 | 304 | GCG | ACG | A73T | Rv1129c | *prpR* |
| MD-051-11 | 1253819 | 0.16 | 335 | GCC | GTC | A293V | Rv1129c | *prpR* |
| MD-052-11 | 1253483 | 0.45 | 322 | GCG | GTG | A351V | Rv1129c | *prpR* |
| MD-046-11 | 1276785 | 0.68 | 26 | AGC | GGC | S322G | Rv1148c |  |
| MD-046-12 | 1276785 | 0.58 | 186 | AGC | GGC | S322G | Rv1148c |  |
| MD-046-14 | 1276785 | 0.62 | 157 | AGC | GGC | S322G | Rv1148c |  |
| MD-048-11 | 1276588 | 0.12 | 373 | GCG | GCC | A387A | Rv1148c |  |
| MD-048-12 | 1276588 | 0.28 | 304 | GCG | GCC | A387A | Rv1148c |  |
| MD-048-13 | 1276588 | 0.32 | 380 | GCG | GCC | A387A | Rv1148c |  |
| MD-048-14 | 1276588 | 0.26 | 172 | GCG | GCC | A387A | Rv1148c |  |
| MD-001-12 | 3717108 | 0.25 | 105 | CCT | GCT | P7A | Rv3331 | *sugI* |
| MD-009-12 | 3717108 | 0.59 | 100 | CCT | ACT | P7T | Rv3331 | *sugI* |
| MD-033-12 | 3717105 | 0.65 | 139 | CAG | TAG | Q6stop | Rv3331 | *sugI* |

## Discussion

Ongoing or recent selection for drug resistance in MTB may be detectable by the presence of unfixed / mixed SNPs in patient isolates [Trauner 2017]. Here we studied sequential isolates from patients receiving treatment for MDR-TB immediately following the publication of the WHO oral MDR-TB treatment guidelines [WHO 2020]. Variation in the use of bedaquiline between the countries in the initial regimen is striking with 100% of the MDR-TB participants recruited in Belarus being initially prescribed bedaquiline but only 15% of participants starting on a regimen containing bedaquiline in Moldova. This was due to the change in the WHO guidelines directly before the study initiation [WHO 2020] substitutions were made for some drugs (notably bedaquiline) in many participants (Table 1). Emphasizing the significant challenge, for countries when initially rolling out changes in recommended therapy, to ensure drug stocks and procurement are in line with treatment recommendations. This is particularly concerning as a regimens “forgiveness for missed doses” may be limited [Garcia-Cremades 2022] and incomplete/nonstandard regimens may provide an opportunity for the rapid development of mycobacterial resistance to new drugs.

Of the five genes that contained mixed SNPs in two or more participants in this study (Table 2) *gyrA* and *pncA* are associated with resistance to quinolones and pyrazinamide respectively (WHO 2020). A literature search for Rv1148c, did not reveal any known link with drug resistance although it has been reported as a recombination hotspot [Phelan 2016]. However, the (Rv1129c, *prp*R) gene has previously been identified in a genome-wide association study as linked with isoniazid resistance (along with other mutations implicated in other first and second-line antibiotics) but no known mechanistic basis for the coincidence of mutations and drug resistance in this gene could be determined [Hicks 2018]. Further work, by the same authors, identified Rv1129c as a transcriptional regulator of propionate metabolism (*prpR*) which conferred tolerance to multiple antibiotics in a macrophage model [Hicks et al. 2018]. A mutation in the Rv1129c was independently reported in a patient who received many years of tuberculosis treatment in Germany [Sonnenkalb 2021]. None of the *prpR* (Rv1129c) mutations observed in this study appear to have been observed in the previous reports [Hicks 2018, Sonnenkalb 2021] but the appearance of mutations in this gene in patients receiving tuberculosis treatment in multiple studies suggests ongoing selection.

Finally, *sugI* mutations have previously been detected in spontaneous *in vitro* d-cycloserine resistance in mutants [Chen 2017] but to our knowledge not from clinical isolates to date. The *sugI* gene encodes a probable sugar-transport integral membrane protein in *M. tuberculosis*. The *sugI* Q6 stop mutation identified in this study (Table 2.) was also detected by Chen *et al*. (2017) in a spontaneous *in vitro* mutant. This loss-of-function mutation was presumed to result in a lower uptake of and thus increased resistance to d-cycloserine [Chen 2017]. Three different mutations in three different participants in our cohort (Table 2) in a gene previously associated with *in vitro* d-cycloserine *in vitro* suggests convergent positive selection likely a consequence of d-cycloserine exposure.

There is extensive knowledge of mutations associated with resistance to the first-line anti-tuberculosis drugs and some of the more established drugs used to treat MDR-TB which was recently summarized in a catalogue of mutation published by the WHO [WHO 2021]. Unfortunately for many of the newer and repurposed drugs resistance associated mutations are much less well studied and clinically relevant resistance mechanisms are yet to be defined. This limits our ability to predict resistance based on WGS data and combined with complex and sometimes lack of fully validated *in vitro* MIC testing is a serious problem, especially for recently introduced drugs [Bateson 2022, Nguyen T.V.A, 2018]. As WGS is increasingly applied to detect resistance mutations and predict susceptibility there is an urgent need to identify and characterize resistance associated associations. The characterization of spontaneous *in vitro* mutants can guide this process but their clinical significance remains uncertain until they are identified in drug resistant clinical isolates [Bergval 2009]. Previous studies that have followed mutation selection during treatment have reported resistance development to individual drugs sometimes involves multiple unfixed mutations [Trauner 2017, Nguyen Q.H. 2018, Nimmo 2020]. This phenomena was also observed in this study, in the *pncA* and Rv0678 genes. Presumably, if these diverse mutants are given the chance to evolve, as a result of continued ineffective treatment, the fittest variants (if any) will become fixed. Thus, the detection of mutations identified previously in *in vitro* spontaneous mutants emerging in clinical isolates exposed to the relevant antibiotic, is of considerable interest, supports an association with *in vivo* resistance, and suggests the drug levels achieved during treatment are insufficiently powerful to block the accumulation of antimicrobial resistance mutants.

### Limitations

We screened for mixed SNPs in short read (Illumina) data from cultured isolates. The analysis of regions of repetitive DNA and duplicated regions in the *M. tuberculosis* genome was not possible. As cultured isolates were sequenced culture bias of mutants and the accumulation of mutations during culture is a possibility. We only screened specific genes of interest for InDels. Also, calling low frequency SNPs remains challenging [Blassel 2021] and as our filtering was stringent in an attempt to avoid overcalling SNPs we may have missed some variability. Others have published *M. tuberculosis* exclusion lists of SNPs that cannot be accurately called at low frequency notably, Marin *et. al* in 2022 published a list of regions that in the exclusion list of regions not accurately called by short read sequencing, 67% of SNPs excluded in our analysis overlap with their excluded regions.

## Conclusion

Although the newly available drugs and regimens have dramatically improved the treatment and outcomes for M(X)DR-TB patients [Ahmad 2018, Nunn 2019, Conradie 2022, Chesov 2021] in patient bacterial adaptation in genes with a reported association with drug resistance was seen in 19 of the 53 patients studied here (supplementary table 3). This was likely at least in part to initially sub-optimal inventory of the newer drugs in one center which resulted in the use of diverse, possibly sub optimal, regimens (Supplementary table 1). If these, or similar mutations, are fixed and accumulate in circulating strains the future effectiveness of these regimens will be undermined, as has happened with previous regimens. Monitoring for emerging mutations, phenotypic resistance, and ensuring adequate supplies of drugs to provide optimal regimens immediately to all eligible patients should be a priority [Nguyen T.V.A 2018]. It is particularly notable that in this cohort of 54 patients the treatment was apparently inadequate to prevent the emergence of d-cycloserine escape mutants, detectable in WGS data, in the second isolate of three patients. We would therefore encourage other groups to screen the *sugI* gene in patients known to have received or receiving d-cycloserine especially for mutations emerging early in the gene (codons 6 and 7).

## Acknowledgements

We would like to acknowledge the support of The Netherlands Organisation of Health, Research and Development (ZonMw) for funding this study (project 541002006 within the antibacterial resistance [ABR] Programme), the help of Alejandra Hernandez Segura for bioinformatics support and preparing R-scrips to facilitate the SNP screening, and the patients who agreed to participate in the study.

## References

Ahmad, N., Ahuja, S. D., Akkerman, O. W., Alffenaar, J. W. C., Anderson, L. F., Baghaei, P., … & Menzies, D. (2018). Treatment correlates of successful outcomes in pulmonary multidrug-resistant tuberculosis: an individual patient data meta-analysis. The Lancet, 392(10150), 821–834.

Anthony, R. M., Tagliani, E., Nikolayevskyy, V., de Zwaan, R., Mulder, A., Kamst, M., … & ERLTB-Net members. (2023). Experiences from 4 Years of Organization of an External Quality Assessment for Mycobacterium tuberculosis Whole-Genome Sequencing in the European Union/European Economic Area. Microbiology Spectrum, 11(1), e02244–22.

Bateson, A., Ortiz Canseco, J., McHugh, T. D., Witney, A. A., Feuerriegel, S., Merker, M., … & Timm, J. (2022). Ancient and recent differences in the intrinsic susceptibility of Mycobacterium tuberculosis complex to pretomanid. Journal of Antimicrobial Chemotherapy, 77(6), 1685–1693.

Bergval, I. L., Schuitema, A. R., Klatser, P. R., & Anthony, R. M. (2009). Resistant mutants of Mycobacterium tuberculosis selected in vitro do not reflect the in vivo mechanism of isoniazid resistance. Journal of Antimicrobial Chemotherapy, 64(3), 515–523.

Castro, R. A., Borrell, S., & Gagneux, S. (2021). The within-host evolution of antimicrobial resistance in Mycobacterium tuberculosis. FEMS microbiology reviews, 45(4), fuaa071.

Luc Blassel, Anna Zhukova, Christian J Villabona-Arenas, Katherine E Atkins, Stéphane Hué, Olivier Gascuel, Drug resistance mutations in HIV: new bioinformatics approaches and challenges, Current Opinion in Virology, Volume 51, 2021, Pages 56–64, ISSN 1879-6257, https://doi.org/10.1016/j.coviro.2021.09.009.

Chen, J., Zhang, S., Cui, P., Shi, W., Zhang, W., & Zhang, Y. (2017). Identification of novel mutations associated with cycloserine resistance in Mycobacterium tuberculosis. Journal of Antimicrobial Chemotherapy, 72(12), 3272–3276.

Chesov, D., Heyckendorf, J., Alexandru, S., Donica, A., Chesov, E., Reimann, M., … & Lange, C. (2021). Impact of bedaquiline on treatment outcomes of multidrug-resistant tuberculosis in a high-burden country. European Respiratory Journal, 57(6).

Coll F, McNerney R, Guerra-Assuncao JA, Glynn JR, Perdigao J, Viveiros M, Portugal I, Pain A, Martin N, Clark TG. 2014. A robust SNP barcode for typing Mycobacterium tuberculosis complex strains. Nature communications 5.

Conradie, F., Bagdasaryan, T. R., Borisov, S., Howell, P., Mikiashvili, L., Ngubane, N., … & Spigelman, M. (2022). Bedaquiline–pretomanid–linezolid regimens for drug-resistant tuberculosis. New England Journal of Medicine, 387(9), 810–823.

Fox, W. S., Strydom, N., Imperial, M. Z., Jarlsberg, L., & Savic, R. M. (2022). Examining nonadherence in the treatment of tuberculosis: The patterns that lead to failure. British Journal of Clinical Pharmacology.

Garcia-Cremades, M., Solans, B. P., Strydom, N., Vrijens, B., Pillai, G. C., Shaffer, C., … & Savic, R. M. (2022). Emerging therapeutics, technologies, and drug development strategies to address patient nonadherence and improve tuberculosis treatment. Annual Review of Pharmacology and Toxicology, 62, 197–210.

Hicks ND, Yang J, Zhang X, Zhao B, Grad YH, Liu L, et al. Clinically prevalent mutations in Mycobacterium tuberculosis alter propionate metabolism and mediate multidrug tolerance. Nat Microbiol. 2018;3:1032–42.

Jajou, R., van der Laan, T., de Zwaan, R., Kamst, M., Mulder, A., de Neeling, A., … & van Soolingen, D. (2019). WGS more accurately predicts susceptibility of Mycobacterium tuberculosis to first-line drugs than phenotypic testing. Journal of Antimicrobial Chemotherapy, 74(9), 2605–2616.

Li, H., Handsaker, B., Wysoker, A., Fennell, T., Ruan, J., Homer, N., … & 1000 Genome Project Data Processing Subgroup. (2009). The Sequence alignment/map (SAM) format and SAMtools. Bioinformatics, 25(16), 2078–2079.

Marin, M., Vargas Jr, R., Harris, M., Jeffrey, B., Epperson, L. E., Durbin, D., … & Farhat, M. R. (2022). Benchmarking the empirical accuracy of short-read sequencing across the M. tuberculosis genome. Bioinformatics, 38(7), 1781–1787.

de Neeling, A. J., Nieboer, L. F. J., Mulder, A., Mariman, R., Anthony, R. M., & van Soolingen, D. (2022). Tracking Mycobacterium tuberculosis sequencing samples using unique spikes of random DNA. Journal of Microbiological Methods, 197, 106482.

Nguyen, Q. H., Contamin, L., Nguyen, T. V. A., & Bañuls, A. L. (2018). Insights into the processes that drive the evolution of drug resistance in Mycobacterium tuberculosis. Evolutionary applications, 11(9), 1498–1511.

Nguyen, T. V. A., Anthony, R. M., Bañuls, A. L., Nguyen, T. V. A., Vu, D. H., & Alffenaar, J. W. C. (2018). Bedaquiline resistance: its emergence, mechanism, and prevention. Clinical Infectious Diseases, 66(10), 1625–1630.

Nimmo, C., Brien, K., Millard, J., Grant, A. D., Padayatchi, N., Pym, A. S., … & Balloux, F. (2020). Dynamics of within-host Mycobacterium tuberculosis diversity and heteroresistance during treatment. EBioMedicine, 55, 102747.

Nunn, A. J., Phillips, P. P., Meredith, S. K., Chiang, C. Y., Conradie, F., Dalai, D., … & Rusen, I. D. (2019). A trial of a shorter regimen for rifampin-resistant tuberculosis. New England Journal of Medicine, 380(13), 1201–1213.

Phelan, J. E., Coll, F., Bergval, I., Anthony, R. M., Warren, R., Sampson, S. L., … & Clark, T. G. (2016). Recombination in pe/ppe genes contributes to genetic variation in Mycobacterium tuberculosis lineages. BMC genomics, 17, 1–12.

Sonnenkalb, L., Strohe, G., Dreyer, V., Andres, S., Hillemann, D., Maurer, F. P., … & Merker, M. (2021). Microevolution of Mycobacterium tuberculosis Subpopulations and Heteroresistance in a Patient Receiving 27 Years of Tuberculosis Treatment in Germany. Antimicrobial agents and chemotherapy, 65(7), e02520–20.

Trauner, A., Liu, Q., Via, L. E., Liu, X., Ruan, X., Liang, L., … & Gao, Q. (2017). The within-host population dynamics of Mycobacterium tuberculosis vary with treatment efficacy. Genome biology, 18(1), 1–17.

Villellas, C., Coeck, N., Meehan, C. J., Lounis, N., de Jong, B., Rigouts, L., & Andries, K. (2017). Unexpected high prevalence of resistance-associated Rv0678 variants in MDR-TB patients without documented prior use of clofazimine or bedaquiline. Journal of Antimicrobial Chemotherapy, 72(3), 684–690.

Wilm, A., Aw, P. P. K., Bertrand, D., Yeo, G. H. T., Ong, S. H., Wong, C. H., … & Nagarajan, N. (2012). LoFreq: a sequence-quality aware, ultra-sensitive variant caller for uncovering cell-population heterogeneity from high-throughput sequencing datasets. Nucleic acids research, 40(22), 11189–11201.

World Health Organization (WHO). WHO Consolidated Guidelines on Tuberculosis Module 4: Treatment Drug-resistant Tuberculosis Treatment. Geneva, WHO, 2020

World Health Organization (WHO), Catalogue of mutations in Mycobacterium tuberculosis complex and their association with drug resistance: supplementary document. supplementary document. 2021, World Health Organization: Geneva

