## Supplementary material for "The appearance of *sugI* mixed loci in three individuals during treatment for MDR-TB, supports the involvement of *sugI* in *Mycobacterium tuberculosis* d-cycloserine resistance *in vivo*": Suplementary Tables

**Suplementary table 1 drugs provided to patinets recruited to the study**

| Country | Participant ID | Intial drugs provided | Drugs provided follow up 1 | Drugs provided follow up 2 | Drugs provided follw up 3 |
|---|---|---|---|---|---|
| Belarus | 1 | lev cyc lin clo bed | lev cyc lin clo bed | lev cyc lin clo bed | lev cyc lin clo bed |
| Belarus | 2 | lev cyc lin clo bed | lev cyc lin clo bed | lev cyc lin clo bed | lev cyc lin clo bed |
| Belarus | 3 | lev cyc lin clo bed | lev cyc lin clo bed | lev cyc lin clo bed | lev cyc lin clo bed |
| Belarus | 4 | lev cyc lin clo bed | lev cyc lin clo bed | lev cyc lin clo bed | lev cyc lin clo bed |
| Belarus | 5 | lev cyc lin clo bed | lev cyc lin clo bed | lev cyc lin clo bed | lev cyc lin clo bed |
| Belarus | 6 | cyc lin clo bed imi amo | cyc lin clo bed imi amo | cyc lin clo bed imi amo | EOS |
| Belarus | 7 | lev cyc lin clo bed | lev cyc lin clo bed | lev cyc lin clo bed | lev cyc lin clo bed |
| Belarus | 8 | lev cyc lin clo bed | lev cyc lin clo bed | lev cyc lin clo bed | lev cyc lin clo bed |
| Belarus | 9 | lev cyc lin clo bed | lev cyc lin clo bed | lev cyc lin clo bed | lev cyc lin clo bed |
| Belarus | 10 | cyc lin clo bed del | cyc lin clo bed del | cyc lin clo bed del | cyc lin clo bed del |
| Belarus | 11 | lev cyc lin clo bed | lev cyc lin clo bed | lev cyc lin clo bed | lev cyc lin clo bed |
| Belarus | 12 | lev cyc lin clo bed | lev cyc lin clo bed | lev cyc lin clo bed | lev cyc lin clo bed |
| Belarus | 13 | lev cyc lin clo bed | lev cyc lin clo bed | lev cyc lin clo bed | lev cyc lin clo bed |
| Belarus | 14 | lev cyc lin clo bed | lev cyc lin clo bed | lev cyc lin clo bed | lev cyc lin clo bed |
| Belarus | 15 | cyc lin clo bed imi amo | cyc lin clo bed imi amo | cyc lin clo bed del | cyc lin clo bed del |
| Belarus | 16 | lin clo bed del | lin clo bed del | lin clo bed del | lin clo bed del |
| Belarus | 17 | lin clo bed del imi amo | lin clo bed del imi amo | lin clo bed del imi amo | lin clo bed del imi amo |
| Belarus | 18 | lev cyc lin clo bed | lev cyc lin clo bed | cyc lin clo bed del | cyc lin clo bed del |
| Belarus | 19 | lev cyc lin clo bed | lev cyc lin clo bed | lev cyc lin clo bed | lev cyc lin clo bed |
| Belarus | 20 | lev cyc lin clo bed | lev cyc lin clo bed | lev cyc lin clo bed | lev cyc lin clo bed |
| Belarus | 21 | lev cyc lin clo bed | lev cyc lin clo bed | EOS | EOS |
| Belarus | 22 | lev cyc lin clo bed | lev cyc lin clo bed | EOS | EOS |
| Belarus | 23 | lev cyc lin clo bed | lev cyc lin clo bed | lev cyc lin clo bed | lev cyc lin clo bed |
| Belarus | 24 | lev cyc lin clo bed | lev cyc lin clo bed | lev cyc lin clo bed | lev cyc lin clo bed |
| Belarus | 25 | lev cyc lin clo bed | lev cyc lin clo bed | lev cyc lin clo bed | lev cyc lin clo bed |
| Belarus | 26 | lev cyc lin clo bed | lev cyc lin clo bed | cyc lin clo bed imi amo | cyc lin clo bed imi amo |
| Belarus | 27 | lev cyc lin clo bed | lev cyc lin clo bed imi amo | cyc lin clo bed imi amo | cyc lin clo bed imi amo |
| Belarus | 28 | lev cyc lin clo bed | lev cyc lin clo bed | lev cyc lin clo bed | lev cyc lin clo bed |
| Belarus | 29 | lev cyc lin clo bed | lev cyc lin clo bed | lev cyc lin clo bed | lev cyc lin clo bed |
| Belarus | 30 | lev cyc lin clo bed | lev cyc lin clo bed | lev cyc lin clo bed | lev cyc lin clo bed |
| Belarus | 31 | lev cyc lin clo bed | lev cyc lin clo bed | lev cyc lin clo bed | lev cyc lin clo bed |
| Moldova | 1 | lev cyc lin bed | lev cyc lin bed | lev cyc lin bed | lev cyc lin bed |
| Moldova | 2 | pza lev ethp cyc lin | pza lev ethp cyc lin | pza lev ethp cyc lin | pza lev ethp cyc lin |
| Moldova | 3 | pza lev ethp cyc lin | pza lev ethp cyc lin pas | pza lev cyc lin bed pas | pza lev cyc lin bed pas |
| Moldova | 4 | pza lev cyc lin bed | pza lev cyc lin bed | pza lev cyc lin | pza lev cyc lin |
| Moldova | 5 | pza lev cyc lin pas | pza lev cyc lin pas | pza lev cyc lin bed | pza cap cyc lin bed |
| Moldova | 6 | pza lev ethp lin clo pas | pza lev ethp lin clo | pza lev lin clo | pza lev lin clo bed |
| Moldova | 7 | pza lev ethp cyc lin pas | pza lev cyc lin pas | EOS | EOS |
| Moldova | 8 | pza cyc lin bed pas imi amo | pza cyc lin bed pas imi amo | pza cyc lin bed pas imi amo | EOS |
| Moldova | 9 | pza lev ethp cyc | pza lev ethp cyc lin pas imi amo | pza lev ethp cyc lin pas imi amo | pza lev cap ethp cyc bed pas |
| Moldova | 10 | pza lev ethp cyc lin pas | pza lev ethp cyc lin pas | pza lev cyc lin | pza lev cap cyc lin imi amo |
| Moldova | 11 | lev cyc lin clo bed | lev cyc lin clo bed | lev cyc lin clo bed | lev cyc lin clo bed |
| Moldova | 12 | lev cyc lin | lev cyc lin | lev cyc lin clo bed | lev cyc lin clo bed |
| Moldova | 13 | lev cyc lin | lev cyc lin | lev cyc lin clo bed | lev cyc lin clo bed |
| Moldova | 14 | pza lev lin pas | pza lev lin pas | pza lev mox lin pas | pza lev mox lin pas |
| Moldova | 15 | pza lev cap ethp cyc lin | pza lev cap ethp cyc lin | pza lev cap ethp cyc lin | pza lev cap ethp cyc lin |
| Moldova | 16 | pza cap cyc lin pas imi amo | pza cap cyc lin pas imi amo | pza cap cyc lin pas imi amo | pza cap cyc lin pas imi amo |
| Moldova | 17 | pza lev cap ethp cyc lin | pza lev cap ethp cyc lin pas | pza lev cap ethp cyc lin pas imi amo | pza lev cap ethp cyc lin bed pas imi amo |
| Moldova | 18 | pza mox cap cyc lin imi amo | pza mox cap cyc lin imi amo | pza mox cap cyc lin bed imi amo | pza cap cyc lin bed pas imi amo |
| Moldova | 19 | lev cyc lin | lev cyc lin | lev cyc lin clo bed | lev cyc lin clo bed |
| Moldova | 20 | lev cyc lin clo bed | lev cyc lin clo bed | lev cyc lin clo bed | EOS |
| Moldova | 21 | pza lev cap cyc lin | pza lev ethp cyc lin imi amo | pza lev ethp cyc lin imi amo | pza lev ethp cyc lin bed imi amo |
| Moldova | 22 | pza lev cap cyc pas imi amo | pza lev cap cyc pas imi amo | pza lev cap cyc pas imi amo | pza lev cap cyc pas imi amo |
| Moldova | 23 | iso rif eth | iso rif eth | iso rif eth | iso rif eth |
| Moldova | 24 | pza lev cap ethp cyc lin pas imi amo | pza lev cap ethp cyc lin pas imi amo | pza lev cap ethp cyc lin pas imi amo | pza lev cap ethp cyc lin bed pas imi amo |
| Moldova | 25 | pza lev cyc lin | pza lev cyc lin | pza lev cyc lin bed | EOS |
| Moldova | 26 | pza cyc lin pas | pza cyc lin pas | pza cyc lin bed pas | pza cyc lin bed pas |
| Moldova | 27 | pza lev cap cyc lin | pza lev cap cyc lin pas imi amo | pza lev cap cyc lin bed pas amo | pza lev cap cyc lin bed pas amo |

| | | | | | |
|---|---|---|---|---|---|
| Moldova | 28 | pza lev cap cyc lin | pza lev cap cyc lin | pza lev cap cyc lin | EOS |
| Moldova | 29 | iso rif eth | iso rif eth | EOS | EOS |
| Moldova | 30 | pza mox cyc lin | pza mox cyc lin | pza mox cyc lin | EOS |
| Moldova | 31 | pza lev ethp cyc lin | pza lev ethp cyc lin | pza lev ethp cyc lin | pza lev ethp cyc lin |
| Moldova | 32 | pza lev cap ethp cyc lin | pza lev cap ethp cyc lin | pza lev cap ethp cyc lin bed | pza lev cap ethp cyc lin bed |
| Moldova | 33 | pza lev cyc lin pas imi amo | pza lev cyc lin pas imi amo | pza lev cyc lin bed pas imi amo | pza lev cyc lin bed pas |
| Moldova | 34 | pza lev cap cyc lin pas imi amo | pza lev cap cyc lin bed pas imi amo | pza lev cap cyc lin bed pas imi amo | pza lev cap cyc lin bed pas |
| Moldova | 35 | pza lev cap ethp cyc lin | pza lev cap ethp cyc lin | pza lev cap ethp cyc lin | pza cap cyc lin |
| Moldova | 36 | pza lev cap cyc lin pas | pza lev cap cyc lin pas | pza lev cap cyc lin pas | pza lev cap cyc lin pas |
| Moldova | 37 | pza lev cap ethp cyc lin pas | pza lev cap ethp cyc lin pas | pza lev cap ethp cyc lin pas | pza lev cap ethp cyc lin pas |
| Moldova | 38 | pza lev cap cyc lin | pza lev cap cyc lin | pza lev cap cyc lin | EOS |
| Moldova | 39 | pza lev cap ethp cyc lin | pza lev cap ethp cyc lin | pza lev cap ethp cyc lin | pza lev cap ethp cyc lin |
| Moldova | 40 | pza cyc lin pas imi amo | pza cyc lin pas imi amo | pza cyc lin pas imi amo | pza cyc lin pas imi amo |
| Moldova | 41 | pza lev cap ethp cyc lin | pza lev cap ethp cyc lin | pza lev cap ethp cyc lin | EOS |
| Moldova | 42 | lev cap ethp cyc lin | lev cap ethp cyc lin | lev cap ethp cyc lin | lev cap ethp cyc lin |
| Moldova | 43 | pza lev cap lin | pza lev cap lin | pza lev cap lin | pza lev cap lin |
| Moldova | 44 | pza lev ethp cyc lin | pza lev ethp cyc lin | pza lev ethp cyc lin | EOS |
| Moldova | 45 | pza lev ethp cyc lin | pza lev ethp cyc lin | pza lev ethp cyc lin | pza lev ethp cyc lin |
| Moldova | 46 | pza cyc lin imi amo | pza cyc lin imi amo | pza cyc lin imi amo | pza cyc lin imi amo |
| Moldova | 47 | pza lev ethp cyc lin | pza lev ethp cyc lin | pza lev ethp cyc lin | EOS |
| Moldova | 48 | pza lev ethp cyc lin imi amo | pza lev ethp cyc lin imi amo | pza lev ethp cyc lin imi amo | pza lev ethp cyc lin imi amo |
| Moldova | 49 | pza ethp cyc lin | pza ethp cyc lin | pza ethp cyc lin | EOS |
| Moldova | 50 | pza cyc lin imi amo | pza cyc lin imi amo | pza cyc lin imi amo | pza cyc lin imi amo |
| Moldova | 51 | pza mox cyc lin pas imi amo | pza mox cyc lin pas imi amo | pza mox cyc lin pas imi amo | pza mox cyc lin pas imi amo |
| Moldova | 52 | pza mox lin imi amo | pza mox lin imi amo | pza mox lin imi amo | pza mox lin imi amo |
| Moldova | 53 | pza lev cyc lin bed | pza lev cyc lin bed | pza lev cyc lin bed | pza lev cyc lin bed |
| Moldova | 54 | pza lev cap cyc lin | pza lev cap cyc lin | pza lev cap cyc lin | pza lev cap cyc lin |
| Moldova | 55 | pza lev cap cyc lin | pza lev cap cyc lin | pza lev cap cyc lin | pza lev cap cyc lin |

**Supplementary table 2**

**List of SNP positions (relative to the reference genome GenBank accession: AL123456.3) excluded from before sreening for confidently mixed SNPs**

| |
|---|
| 71 |
| 39066 |
| 55553 |
| 71584 |
| 84832 |
| 334751 |
| 335720 |
| 335885 |
| 335906 |
| 336504 |
| 336728 |
| 338648 |
| 338669 |
| 338719 |
| 338810 |
| 338876 |
| 338903 |
| 580772 |
| 631392 |
| 832109 |
| 839309 |
| 839463 |
| 839534 |
| 839821 |
| 840272 |
| 840329 |
| 840447 |
| 840496 |
| 840515 |
| 888774 |

| |
|---|
| 916690 |
| 964636 |
| 964666 |
| 964801 |
| 1067665 |
| 1093640 |
| 1093928 |
| 1094375 |
| 1096205 |
| 1096235 |
| 1190093 |
| 1191497 |
| 1191741 |
| 1340578 |
| 1342096 |
| 1443428 |
| 1480972 |
| 1481337 |
| 1612648 |
| 1644309 |
| 1644362 |
| 1789446 |
| 1789516 |
| 1789650 |
| 1983235 |
| 2030848 |
| 2133475 |
| 2163375 |
| 2163444 |
| 2163493 |
| 2165503 |
| 2165902 |
| 2165938 |
| 2196715 |
| 2262026 |
| 2266442 |
| 2266487 |
| 2266583 |
| 2266624 |
| 2296181 |
| 2306306 |
| 2338457 |
| 2339605 |
| 2339907 |
| 2372460 |
| 2372517 |
| 2372550 |
| 2401825 |
| 2439401 |
| 2439519 |
| 2531964 |
| 2591829 |
| 2592310 |
| 2631599 |
| 2631620 |
| 2631779 |
| 2631827 |
| 2632922 |
| 2635482 |
| 2635496 |
| 2635512 |

2638997
2704886
2751675
2829294
2866489
2866503
2866551
2866956
2945101
2983613
3007146
3123477
3131473
3232703
3232759
3232815
3336646
3377198
3378095
3379432
3380534
3391069
3477900
3478272
3478467
3478698
3478719
3482717
3528102
3529067
3537806
3558767
3590689
3730466
3730519
3730616
3730642
3730648
3730684
3730993
3732247
3732310
3732344
3732553
3735865
3736024
3738521
3820407
3820429
3820486
3820523
3820545
3842452
3843559
3846622
3846687
3846764
3847022
3847039
3847052
3847073

3847112
3847153
3847511
3847528
3847659
3883711
3928299
3929084
3930476
3932564
3932579
3934699
3941515
3943705
3947248
3948712
4053050
4060100
4060230
4120926
4120983
4359135
4359165
4359183

**Suplementary file 3**
**Sequence files available (NCBI bioproject PRJNA825716) / reason for exclusion from the analysis**

| | ID (country-participan | Patient count per site | Coll* classification / reasn for exclusion |
|---|---|---|---|
| 1 | BE-001-06 | 1 | 2.2.1 |
| 2 | BE-001-07 | | 2.2.1 |
| 3 | BE-002-06 | 2 | 4.8 |
| 4 | BE-002-07 | | 4.8 |
| 5 | BE-003-06 | 3 | 4.3.3 |
| 6 | BE-004-6 | 4 | 2.2.1 |
| 7 | BE-004-7 | | 2.2.1 |
| 8 | BE-004-14 | | 2.2.1 |
| 9 | BE-005-6 | 5 | 2.2.1 |
| 10 | BE-005-7 | | 2.2.1 |
| 11 | BE-006-6 | 6 | 2.2.1 |
| 12 | BE-006-7 | | 2.2.1 |
| 13 | BE-006-8 | | 2.2.1 |
| 14 | BE-006-9 | | 2.2.1 |
| 15 | BE-007-6 | 7 | 2.2.1 |
| 16 | BE-008-6 | 8 | 4.3.3 |
| 17 | BE-008-7 | | 4.3.3 |
| 18 | BE-0010-6 | 9 | 2.2.1 |
| 19 | BE-0010-7 | | 2.2.1 |
| 20 | BE-0010-8 | | 2.2.1 |
| 21 | BE-0011-6 | 10 | 4.8 |
| 22 | BE-0011-7 | | 4.8 |
| 23 | BE-0011-8 | | 4.8 |
| 24 | BE-0012-6 | 11 | 2.2.1 |
| 25 | BE-0012-7 | | 2.2.1 |
| 26 | MD-001-11 | 1 | 2.2.1 |
| 27 | MD-001-12 | | 2.2.1 |
| 28 | MD-001-13 | | 2.2.1 |
| 29 | MD-002-11 | 2 | 4.3.3 |
| 30 | MD-002-14 | | 4.3.3 |
| 31 | MD-003-11 | 3 | 4.2.1 |
| 32 | MD-003-12 | | 4.2.1 |
| 33 | MD-003-13 | | 4.2.1 |
| 34 | MD-005-11 | 4 | 2.2.1 |

| | | | |
|---|---|---|---|
| 35 | MD-005-12 | | 2.2.1 |
| 36 | MD-005-13 | | 2.2.1 |
| 37 | MD-007-11 | 5 | 4.2.1 |
| 38 | MD-007-12 | | 4.2.1 |
| 39 | MD-008-12 | 6 | 2.2.1 |
| 40 | MD-009-11 | 7 | 2.2.1 |
| 41 | MD-009-12 | | 2.2.1 |
| 42 | MD-009-13 | | 2.2.1 |
| 43 | MD-010-11 | 8 | 4.2.1 |
| 44 | MD-010-12 | | 4.2.1 |
| 45 | MD-010-13 | | 4.2.1 |
| 46 | MD-011-11 | 9 | 4.2.1 |
| 47 | MD-011-12 | | 4.2.1 |
| 48 | MD-011-13 | | 4.2.1 |
| 49 | MD-013-11 | 10 | 2.2.1 |
| 50 | MD-015-11 | 11 | 2.2.1 |
| 51 | MD-016-11 | 12 | 2.2.1 |
| 52 | MD-016-12 | | 2.2.1 |
| 53 | MD-017-11 | 13 | 2.2.1 |
| 54 | MD-017-12 | | MIXED genotype excluded |
| 55 | MD-017-13 | | 2.2.1 |
| 56 | MD-018-12 | 14 | 2.2.1 |
| 57 | MD-020-11 | 15 | 4.2.1 |
| 58 | MD-021-11 | 16 | 2.2.1 |
| 59 | MD-021-12 | | 2.2.1 |
| 60 | MD-022-11 | 17 | 2.2.1 |
| 61 | MD-022-12 | | 2.2.1 |
| 62 | MD-023-11 | 18 | 4.2.1 not treated as MDR |
| 63 | MD-024-11 | 19 | 4.2.1 |
| 64 | MD-024-12 | | 4.2.1 |
| 65 | MD-025-11 | 20 | 2.2.1 |
| 66 | MD-025-12 | | 2.2.1 |
| 67 | MD-025-13 | | 2.2.1 |
| 68 | MD-025-14 | | 2.2.1 |
| 69 | MD-026-12 | 21 | 2.2.1 |
| 70 | MD-026-13 | | 2.2.1 |
| 71 | MD-027-11 | 22 | 2.2.1 |
| 72 | MD-027-12 | | 2.2.1 |
| 73 | MD-027-14 | | 4.2.1 different genotype to first isolate(s) |
| 74 | MD-029-11 | 23 | 4.3.3 not treated as MDR |
| 75 | MD-032-11 | 24 | 2.2.1 |
| 76 | MD-032-12 | | 2.2.1 |
| 77 | MD-033-11 | 25 | 2.2.1 |
| 78 | MD-033-12 | | 2.2.1 |
| 79 | MD-034-11 | 26 | 4.2.1 |
| 80 | MD-034-12 | | 4.2.1 |
| 81 | MD-035-11 | 27 | 2.2.1 |
| 82 | MD-035-12 | | 2.2.1 |
| 83 | MD-035-13 | | 2.2.1 |
| 84 | MD-036-12 | 28 | 2.2.1 |
| 85 | MD-037-11 | 29 | 4.2.1 |
| 86 | MD-038-11 | 30 | 2.2.1 |
| 87 | MD-039-11 | 31 | 4.2.1 |
| 88 | MD-039-12 | | 4.2.1 |
| 89 | MD-039-13 | | 4.2.1 |
| 90 | MD-040-11 | 32 | 4.2.1 |
| 91 | MD-040-12 | | 4.2.1 |
| 92 | MD-040-13 | | 4.2.1 |
| 93 | MD-040-14 | | 4.2.1 |
| 94 | MD-041-11 | 33 | 4.2.1 |
| 95 | MD-041-12 | | 4.2.1 |
| 96 | MD-042-11 | 34 | 4.2.1 |
| 97 | MD-042-12 | | 4.2.1 not closely related to previous isolate from the same individual |
| 98 | MD-043-13 | 35 | 4.2.1 |
| 99 | MD-045-11 | 36 | 2.2.1 |
| 100 | MD-045-12 | | 2.2.1 |
| 101 | MD-046-11 | 37 | 2.2.1 |
| 102 | MD-046-12 | | 2.2.1 |
| 103 | MD-046-13 | | 2.2.1 not closely related to other isolates from the same individual |
| 104 | MD-046-14 | | 2.2.1 |
| 105 | MD-047-11 | 38 | 4.1.2.1 |

| | | | |
|---|---|---|---|
| 106 | MD-047-12 | | 4.1.2.1 |
| 107 | MD-048-11 | 39 | 4.2.1 |
| 108 | MD-048-12 | | 4.2.1 |
| 109 | MD-048-13 | | 4.2.1 |
| 110 | MD-048-14 | | 4.2.1 |
| 111 | MD-050-11 | 40 | 2.2.1 |
| 112 | MD-050-12 | | 2.2.1 |
| 113 | MD-051-11 | 41 | 4.2.1 |
| 114 | MD-051-12 | | 4.2.1 |
| 115 | MD-051-13 | | 4.2.1 |
| 116 | MD-052-11 | 42 | 4.2.1 |
| 117 | MD-052-12 | | 4.2.1 |
| 118 | MD-052-13 | | 4.2.1 not closely related to previous isolate from the same individual |
| 119 | MD-053-11 | 43 | 4.2.1 |
| 120 | MD-053-12 | | 4.2.1 |
| 121 | MD-053-13 | | 4.2.1 |
| 122 | MD-054-11 | 44 | 2.2.1 |
| 123 | MD-054-12 | | 2.2.1 |
| 124 | MD-054-13 | | 2.2.1 |
| 125 | MD-055-11 | 45 | 4.2.1 |

* Coll, F., McNerney, R., Guerra-Assunção, J. A., Glynn, J. R., Perdigão, J., Viveiros, M., ... & Clark, T. G. (2014). A robust SNP barcode for typing Mycobacterium tuberculosis complex strains. Nature communications, 5(1), 4812.

**Suplementary table 4**

**Position and frequency of the confident mixed SNPs identifed (in serial isolates)**

| Site | Sample | Postion | Quality | | | Proprtion mutant SNP | Base called froward, reverse | Codon Wild type | Codon mutant | | AA change | Gene | Gene name |
|---|---|---|---|---|---|---|---|---|---|---|---|---|---|
| Belarus | 01-07 | 1258921 | 7760 | 318 | | 0.77673 | 30,39,111,136 | CAT | TAT | 0 | H533Y | Rv1132 | . |
| Belarus | 01-06 | 777262 | 3583 | 302 | | 0.430464 | 85,87,62,68 | ACC | CCC | 0 | T407P | Rv0676c | mmpL5 |
| Belarus | 01-07 | 2625924 | 1120 | 264 | | 0.253788 | 117,80,39,28 | GCA | GCG | 2 | A83A | Rv2346c | esxO |
| Belarus | 02-06 | 2296275 | 870 | 407 | | 0.589681 | 59108117123 | CAC | CGC | 1 | H3571R | Rv2048c | pks12 |
| Belarus | 02-07 | 2296275 | 773 | 495 | | 0.60404 | 67128154145 | CAC | CGC | 1 | H3571R | Rv2048c | pks12 |
| Belarus | 02-06 | 4089777 | 9570 | 408 | | 0.737745 | 41,66,134,167 | ACG | CCG | 0 | T333P | Rv3649 | Rv3649 |
| Belarus | 02-07 | 4089777 | 11742 | 467 | | 0.768737 | 44,64,161,198 | ACG | CCG | 0 | T333P | Rv3649 | Rv3649 |
| Belarus | 02-06 | 653158 | 1138 | 484 | | 0.117769 | 198,228,29,28 | GCC | ACC | 0 | A130T | Rv0562 | grcC1 |
| Belarus | 02-06 | 4138521 | 2985 | 408 | | 0.306373 | 170,111,71,54 | ATG | ACG | 1 | M412T | Rv3696c | glpK |
| Belarus | 03-03 | 1276588 | 1509 | 566 | | 0.150177 | 266,215,68,17 | GCG | GCC | 2 | A387A | Rv1148c | Rv1148c |
| Belarus | 4-06 | 796289 | 2755 | 330 | | 0.318182 | 135,90,66,39 | GAT | GAC | 2 | D257D | Rv0696 | Rv0696 |
| Belarus | 4-07 | 796289 | 2950 | 274 | | 0.386861 | 101,67,63,43 | GAT | GAC | 2 | D257D | Rv0696 | Rv0696 |
| Belarus | 4-08 | 796289 | 10886 | 304 | | 1 | 0,0,188,116 | GAT | GAC | 2 | D257D | Rv0696 | Rv0696 |
| Belarus | 4-06 | 3277158 | 1230 | 336 | | 0.154762 | 152,132,24,28 | GCG | GTG | 1 | A1853V | Rv2940c | mas |
| Belarus | 05-7 | 620371 | 1162 | 368 | | 0.144022 | 155,160,24,29 | GCC | ACC | 0 | A161T | Rv0529 | ccsA |
| Belarus | 06-7 | 536438 | 273 | 189 | | 0.079365 | 70,104,8,7 | GTG | TTG | 0 | V24L | Rv0447c | ufaA1 |
| Belarus | 06-08 | 536438 | 6063 | 184 | | 0.98913 | 2,0,65,117 | GTG | TTG | 0 | V24L | Rv0447c | ufaA1 |
| Belarus | 06-09 | 536438 | 375 | 163 | | 0.122699 | 50,93,4,16 | GTG | TTG | 0 | V24L | Rv0447c | ufaA1 |
| Belarus | 06-6 | 619039 | 1676 | 208 | | 0.307692 | 62,82,27,37 | CAG | CAT | 2 | Q245H | Rv0528 | Rv0528 |
| Belarus | 06-7 | 1937288 | 617 | 310 | | 0.096774 | 172,108,16,14 | CGC | CAC | 1 | R310H | Rv1708 | Rv1708 |
| Belarus | 06-08 | 1937288 | 5303 | 283 | | 0.60424 | 63,48,97,74 | CGC | CAC | 1 | R310H | Rv1708 | Rv1708 |
| Belarus | 06-7 | 3101510 | 2645 | 292 | | 0.332192 | 103,92,45,52 | GAG | GAC | 2 | E24D | Rv2791c | Rv2791c |
| Belarus | 06-7 | 3615605 | 1755 | 264 | | 0.257576 | 110,86,34,34 | GCG | TCG | 0 | A667S | Rv3239c | Rv3239c |
| Belarus | 08-06 | 207399 | 2235 | 275 | | 0.316364 | 81,107,35,52 | ATG | GTG | 0 | M196V | Rv0175 | Rv0175 |
| Belarus | 08-07 | 207399 | 2842 | 298 | | 0.372483 | 92,95,53,58 | ATG | GTG | 0 | M196V | Rv0175 | Rv0175 |
| Belarus | 08-06 | 495753 | 5392 | 264 | | 0.666667 | 42,46,79,97 | GCC | GTC | 1 | A521V | Rv0410c | pknG |
| Belarus | 08-07 | 495753 | 5008 | 305 | | 0.557377 | 68,67,78,92 | GCC | GTC | 1 | A521V | Rv0410c | pknG |
| Belarus | 08-07 | 276505 | 3685 | 369 | | 0.371274 | 137,95,84,53 | CTT | TTT | 0 | L150F | Rv0231 | fadE4 |
| Belarus | 08-06 | 385176 | 1709 | 319 | | 0.213166 | 113,138,32,36 | GCC | GCT | 2 | A256A | Rv0317c | glpQ2 |
| Belarus | 010-06 | 437274 | 7565 | 267 | | 0.850187 | 20,19,105,122 | ACC | GCC | 0 | T139A | Rv0358 | Rv0358 |
| Belarus | 010-07 | 437274 | 9082 | 261 | | 0.992337 | 1,1,125,134 | ACC | GCC | 0 | T139A | Rv0358 | Rv0358 |
| Belarus | 010-08 | 437274 | 798 | 249 | | 0.148594 | 104,107,18,19 | ACC | GCC | 0 | T139A | Rv0358 | Rv0358 |
| Belarus | 010-06 | 600096 | 7074 | 267 | | 0.827715 | 22,24,113,108 | GCC | GCG | 2 | A966A | Rv0507 | mmpL2 |
| Belarus | 010-07 | 600096 | 8484 | 252 | | 0.984127 | 2,2,127,121 | GCC | GCG | 2 | A966A | Rv0507 | mmpL2 |
| Belarus | 010-08 | 600096 | 731 | 269 | | 0.130112 | 122,112,12,23 | GCC | GCG | 2 | A966A | Rv0507 | mmpL2 |
| Belarus | 010-06 | 1622400 | 1081 | 334 | | 0.155689 | 118,163,24,28 | ATG | ACG | 1 | M98T | Rv1443c | Rv1443c |

| | | | | | | | | | | | | | |
|---|---|---|---|---|---|---|---|---|---|---|---|---|---|
| Belarus | 010-08 | 1622400 | 7665 | 268 | 0.850746 | 17,23,91,137 | ATG | ACG | | 1 | M98T | Rv1443c | Rv1443c |
| Belarus | 010-06 | 3525768 | 8068 | 285 | 0.852632 | 21,21,130,113 | CGG | CTG | | 1 | R546L | Rv3157 | nuoM |
| Belarus | 010-07 | 3525768 | 9335 | 264 | 0.988636 | 1,2,140,121 | CGG | CTG | | 1 | R546L | Rv3157 | nuoM |
| Belarus | 010-08 | 3525768 | 811 | 323 | 0.114551 | 144,142,16,21 | CGG | CTG | | 1 | R546L | Rv3157 | nuoM |
| Belarus | 010-07 | 3597979 | 6319 | 307 | 0.651466 | 54,53,106,94 | . | . | . | | . | . | . |
| Belarus | 010-06 | 4020249 | 8929 | 332 | 0.849398 | 26,24,148,134 | ATC | ATA | | 2 | I36I | Rv3578 | arsB2 |
| Belarus | 010-07 | 4020249 | 11250 | 335 | 0.99403 | 1,1,178,155 | ATC | ATA | | 2 | I36I | Rv3578 | arsB2 |
| Belarus | 010-08 | 4020249 | 1189 | 326 | 0.159509 | 153,121,23,29 | ATC | ATA | | 2 | I36I | Rv3578 | arsB2 |
| Belarus | 11-06 | 957390 | 3374 | 246 | 0.479675 | 66,62,64,54 | GGC | GGA | | 2 | G366G | Rv0860 | fadB |
| Belarus | 11-07 | 957390 | 3460 | 286 | 0.433566 | 81,81,70,54 | GGC | GGA | | 2 | G366G | Rv0860 | fadB |
| Belarus | 11-08 | 957390 | 1141 | 302 | 0.175497 | 119,130,23,30 | GGC | GGA | | 2 | G366G | Rv0860 | fadB |
| Belarus | 11-07 | 1278583 | 174 | 317 | 0.605678 | 48,77,108,84 | . | . | . | | . | Rv1150 | Rv1150 |
| Belarus | 11-08 | 1278583 | 515 | 432 | 0.643519 | 57,97,174,104 | . | . | . | | . | Rv1150 | Rv1150 |
| Belarus | 11-08 | 761155 | 3326 | 374 | 0.336898 | 119,129,73,53 | TCG | TTG | | 1 | S450L | Rv0667 | rpoB |
| Belarus | 012-06 | 1490322 | 4082 | 281 | 0.487544 | 64,80,56,81 | GGT | GAT | | 1 | G664D | Rv1326c | glgB |
| Belarus | 012-06 | 1495989 | 1474 | 287 | 0.229965 | 127,93,26,40 | CTG | TTG | | 0 | L476L | Rv1328 | glgP |
| Belarus | 012-07 | 1495989 | 2489 | 261 | 0.367816 | 86,79,54,42 | CTG | TTG | | 0 | L476L | Rv1328 | glgP |
| Belarus | 012-06 | 1533539 | 481 | 308 | 0.155844 | 116,144,27,21 | TGG | TCG | | 1 | W32S | Rv1361c | PPE19 |
| Belarus | 012-07 | 1533539 | 3040 | 295 | 0.450847 | 85,77,71,62 | TGG | TCG | | 1 | W32S | Rv1361c | PPE19 |
| Moldova | 1-11 | 950597 | 2620 | 420 | 0.266667 | 159,147,60,52 | ATA | ATG | | 2 | I174M | Rv0853c | pdc |
| Moldova | 1-11 | 2651742 | 2351 | 335 | 0.283582 | 139,100,54,41 | . | . | . | | . | . | na |
| Moldova | 1-11 | 4338596 | 1632 | 496 | 0.6875 | 0,0,150,191 | . | . | . | | . | . | na |
| Moldova | 1-12 | 3717108 | 649 | 106 | 0.254717 | 33,46,12,15 | CCT | GCT | | 0 | P7A | Rv3331 | sugI |
| Moldova | 1-12 | 4338596 | 589 | 203 | 0.689655 | 0,0,67,73 | . | . | . | | . | . | na |
| Moldova | 1-13 | 1202298 | 2864 | 110 | 0.818182 | 11,9,39,51 | AAA | AAC | | 2 | K194N | Rv1077 | cbs |
| Moldova | 1-13 | 3994300 | 4276 | 148 | 0.864865 | 10,10,54,74 | AGT | ACT | | 1 | S539T | Rv3554 | fdxB |
| Moldova | 1-13 | 4338596 | 378 | 144 | 0.6875 | 0,0,45,54 | . | . | . | | . | . | na |
| Moldova | 2-11 | 2721013 | 1073 | 154 | 0.318182 | 49,55,28,21 | CGT | CGC | | 2 | R255R | Rv2424c | Rv2424c |
| Moldova | 2-14 | 561081 | 6166 | 450 | 0.493333 | 119,109,128,94 | NA | GTG | | 211 | V211L | 0 | Rv0470c |
| Moldova | 2-14 | 1278583 | 509 | 422 | 0.677725 | 49,87,168,118 | NA | . | . | | . | 0 | Rv1150 |
| Moldova | 2-14 | 2721013 | 2945 | 416 | 0.334135 | 145,132,66,73 | NA | CGT | | 255 | R255R | 0 | Rv2424c |
| Moldova | 3-11 | 3523862 | 1139 | 350 | 0.14 | 167,134,34,15 | GTG | GTT | | 2 | V543V | Rv3156 | nuoL |
| Moldova | 3-11 | 4162514 | . | 504 | 0.31746 | 148,196,62,98 | ACC | AGC | | 1 | T70S | Rv3719 | Rv3719 |
| Moldova | 3-12 | 3058572 | 1279 | 133 | 0.353383 | 37,49,24,23 | . | . | . | | . | . | na |
| Moldova | 3-13 | 2436573 | 3253 | 116 | 0.853448 | 9,8,50,49 | CGC | CAC | | 1 | R222H | Rv2174 | mptA |
| Moldova | 3-13 | 4142697 | 1354 | 120 | 0.425 | 35,34,25,26 | GGT | GGG | | 2 | G218G | Rv3699 | Rv3699 |
| Moldova | 5-11 | 3942440 | 2305 | 106 | 0.726415 | 16,10,44,33 | . | . | . | | . | . | . |
| Moldova | 5-12 | 794821 | 4569 | 161 | 0.84472 | 9,16,61,75 | TCC | TTC | | 1 | S36F | Rv0695 | Rv0695 |
| Moldova | 5-12 | 3942440 | 271 | 42 | 0.238095 | 18,14,6,4 | . | . | . | | . | . | na |
| Moldova | 5-13 | 3942440 | 1473 | 42 | 1 | 0,0,22,20 | . | . | . | | . | . | na |
| Moldova | 5-13 | 4392648 | 634 | 120 | 0.208333 | 37,58,15,10 | GTC | GTA | | 2 | V142V | Rv3907c | pcnA |
| Moldova | 9-11 | 192856 | 6970 | 321 | 0.676012 | 59,45,113,104 | GTC | ATC | | 0 | V94I | Rv0162c | adhE1 |
| Moldova | 9-12 | 3717108 | 1843 | 100 | 0.59 | 16,25,22,37 | CCT | ACT | | 0 | P7T | Rv3331 | sugI |
| Moldova | 10-13 | 3585370 | 580 | 81 | 0.271605 | 21,38,12,10 | TTC | CTC | | 0 | F123L | Rv3208 | Rv3208 |
| Moldova | 11-11 | 2195799 | 2669 | 338 | 0.298817 | 102,135,38,63 | CCC | TCC | | 0 | P46S | Rv1944c | Rv1944c |
| Moldova | 11-12 | 2195799 | 2855 | 155 | 0.606452 | 23,38,39,55 | CCC | TCC | | 0 | P46S | Rv1944c | Rv1944c |
| Moldova | 11-13 | 1719468 | 1163 | 157 | 0.286624 | 46,65,19,26 | CCG | CAG | | 1 | P248Q | Rv1524 | Rv1524 |
| Moldova | 11-13 | 2195799 | 2118 | 171 | 0.450292 | 33,61,28,49 | CCC | TCC | | 0 | P46S | Rv1944c | Rv1944c |
| Moldova | 11-13 | 8416 | 1763 | 316 | 0.224684 | 98,146,24,47 | GTC | GCC | | 1 | V372A | Rv0006 | gyrA |
| Moldova | 13-11 | 1913092 | 3191 | 407 | 0.326781 | 139,134,56,77 | GGC | GGA | | 2 | G38G | Rv1688 | mpg |
| Moldova | 13-11 | 3796796 | 3194 | 394 | 0.314721 | 133,137,60,64 | ATC | ATT | | 2 | I214I | Rv3382c | Rv3382c |
| Moldova | 15-11 | 852537 | 13507 | 456 | 0.881579 | 23,30,177,225 | CGG | TGG | | 0 | R48W | Rv0758 | phoR |
| Moldova | 16-12 | 27695 | 2063 | 123 | 0.585366 | 27,24,35,37 | GGC | GTC | | 1 | G34V | Rv0023 | Rv0023 |
| Moldova | 16-12 | 157166 | 2003 | 136 | 0.492647 | 31,36,35,32 | CTT | CTC | | 2 | L145L | Rv0129c | fbpC |
| Moldova | 16-12 | 238256 | 2106 | 112 | 0.607143 | 24,20,37,31 | ACC | ATC | | 1 | T47I | Rv0201c | Rv0201c |
| Moldova | 16-12 | 656061 | 2023 | 169 | 0.426035 | 48,49,35,37 | ATC | ATA | | 2 | I470I | Rv0565c | Rv0565c |
| Moldova | 16-12 | 778994 | 1149 | 119 | 0.344538 | 46,32,23,18 | AGC | ATC | | 1 | S2I | Rv0678 | Rv0678 |
| Moldova | 16-12 | 1131480 | 1422 | 96 | 0.489583 | 22,27,27,20 | . | . | . | | . | . | . |
| Moldova | 16-12 | 1442746 | 1316 | 82 | 0.536585 | 16,22,19,25 | . | . | . | | . | . | . |
| Moldova | 16-12 | 2288920 | 2364 | 133 | 0.56391 | 33,25,37,38 | GGA | CGA | | 0 | G108R | Rv2043c | pncA |
| Moldova | 16-12 | 2289081 | 1616 | 119 | 0.470588 | 34,29,31,25 | CCG | CTG | | 1 | P54L | Rv2043c | pncA |

| | | | | | | | | | | | | | |
|---|---|---|---|---|---|---|---|---|---|---|---|---|---|
| Moldova | 16-12 | 3799672 | 527 | 69 | 0.304348 | 20,28,10,11 | TCG | TTG | | 1 | S91L | Rv3385c | vapB46 |
| Moldova | 16-11 | 27695 | 13613 | 469 | 0.886994 | 30,23,218,198 | GGC | GTC | | 1 | G34V | Rv0023 | Rv0023 |
| Moldova | 16-11 | 157166 | 447 | 477 | 0.060797 | 194,254,11,18 | CTT | CTC | | 2 | L145L | Rv0129c | fbpC |
| Moldova | 16-11 | 238256 | 13738 | 437 | 0.913043 | 21,17,219,180 | ACC | ATC | | 1 | T47I | Rv0201c | Rv0201c |
| Moldova | 16-11 | 656061 | 537 | 454 | 0.059471 | 186,241,11,16 | ATC | ATA | | 2 | I470I | Rv0565c | Rv0565c |
| Moldova | 16-11 | 778994 | 745 | 335 | 0.101493 | 172,129,19,15 | AGC | ATC | | 1 | S2I | Rv0678 | Rv0678 |
| Moldova | 16-11 | 1131480 | 9486 | 322 | 0.881988 | 19,18,149,135 | . | . | . | | . | . | na |
| Moldova | 16-11 | 1442746 | 223 | 349 | 0.054441 | 154,174,6,13 | . | . | . | | . | . | na |
| Moldova | 16-11 | 2288920 | 10567 | 419 | 0.761337 | 43,57,148,171 | GGA | CGA | | 0 | G108R | Rv2043c | pncA |
| Moldova | 16-11 | 2289081 | 468 | 412 | 0.058252 | 224,164,10,14 | CCG | CTG | | 1 | P54L | Rv2043c | pncA |
| Moldova | 16-11 | 3799672 | 656 | 345 | 0.092754 | 146,167,15,17 | TCG | TTG | | 1 | S91L | Rv3385c | vapB46 |
| Moldova | 21-11 | 3805103 | 1570 | 371 | 0.188679 | 145,155,35,35 | AGA | ACA | | 1 | R80T | Rv3390 | lpqD |
| Moldova | 22-11 | 884449 | 1506 | 136 | 0.389706 | 47,36,32,21 | TCC | CCC | | 0 | S118P | Rv0790c | Rv0790c |
| Moldova | 22-12 | 596957 | 1382 | 208 | 0.264423 | 72,81,32,23 | GGC | AGC | | 0 | G67S | Rv0506 | mmpS2 |
| Moldova | 22-12 | 884449 | 2362 | 140 | 0.55 | 40,23,48,29 | TCC | CCC | | 0 | S118P | Rv0790c | Rv0790c |
| Moldova | 22-12 | 2283389 | 810 | 123 | 0.243902 | 49,44,15,15 | CTG | CTT | | 2 | L111L | Rv2037c | Rv2037c |
| Moldova | 22-12 | 2539414 | 693 | 149 | 0.208054 | 63,54,19,12 | ACC | GCC | | 0 | T239A | Rv2265 | Rv2265 |
| Moldova | 22-12 | 3597009 | 737 | 179 | 0.173184 | 79,69,11,20 | CGC | TGC | | 0 | R176C | Rv3220c | Rv3220c |
| Moldova | 23-11 | 1096470 | 7413 | 644 | 0.492236 | 1.40187E+11 | . | . | . | | . | . | na |
| Moldova | 24-12 | 4168058 | 1418 | 81 | 0.567901 | 11,24,19,27 | GCG | GTG | | 1 | A24V | Rv3722c | Rv3722c |
| Moldova | 24-11 | 4168058 | 14019 | 471 | 0.876858 | 24,34,169,244 | GCG | GTG | | 1 | A24V | Rv3722c | Rv3722c |
| Moldova | 25-14 | 1147810 | 1320 | 179 | 0.290503 | 73,53,33,19 | CAG | CAA | | 2 | Q264Q | Rv1026 | Rv1026 |
| Moldova | 26-12 | 2054204 | 1699 | 408 | 0.171569 | 164,174,33,37 | GCA | CCA | | 0 | A386P | Rv1812c | Rv1812c |
| Moldova | 26-12 | 2095631 | 3927 | 352 | 0.397727 | 101,111,70,70 | GTC | CTC | | 0 | V180L | Rv1845c | blaR |
| Moldova | 26-12 | 4309804 | 4862 | 253 | 0.616601 | 43,54,70,86 | ATG | ACG | | 1 | M253T | Rv3835 | Rv3835 |
| Moldova | 26-13 | 900176 | 3592 | 150 | 0.74 | 18,21,52,59 | CGT | CGC | | 2 | R385R | Rv0806c | cpsY |
| Moldova | 26-13 | 1607213 | 5355 | 193 | 0.829016 | 18,14,84,76 | GAC | GAT | | 2 | D276D | Rv1430 | PE16 |
| Moldova | 26-13 | 2054204 | 5527 | 169 | 0.934911 | 6,3,76,82 | GCA | CCA | | 0 | A386P | Rv1812c | Rv1812c |
| Moldova | 26-13 | 4181263 | 3732 | 133 | 0.834586 | 9,13,51,60 | ACG | ATG | | 1 | T153M | Rv3730c | Rv3730c |
| Moldova | 26-13 | 4309804 | 3195 | 92 | 0.967391 | 1,2,32,57 | ATG | ACG | | 1 | M253T | Rv3835 | Rv3835 |
| Moldova | 29-11 | 2626149 | 8545 | 373 | 0.764075 | 45,43,131,154 | GGT | GGG | | 2 | G8G | Rv2346c | esxO |
| Moldova | 29-11 | 3086769 | 1743 | 475 | 0.16 | 167,232,35,41 | . | . | . | | . | . | na |
| Moldova | 29-11 | 3342824 | 11387 | 461 | 0.770065 | 39,67,137,218 | GAA | GAC | | 2 | E220D | Rv2985 | mutT1 |
| Moldova | 29-11 | 4200213 | 10381 | 513 | 0.649123 | 82,97,153,180 | AGA | AGG | | 2 | R3R | Rv3753c | Rv3753c |
| Moldova | 32-11 | 482558 | 14110 | 455 | 0.896703 | 22,25,213,195 | GTC | GGC | | 1 | V225G | Rv0402c | mmpL1 |
| Moldova | 32-11 | 1110397 | 7620 | 507 | 0.514793 | 1.05141E+11 | ATG | ATA | | 2 | M43I | Rv0994 | moeA1 |
| Moldova | 32-11 | 4074497 | 16501 | 525 | 0.910476 | 28,19,273,205 | ATG | ATC | | 2 | M288I | Rv3635 | Rv3635 |
| Moldova | 32-12 | 482558 | 6954 | 197 | 0.994924 | 1,0,94,102 | GTC | GGC | | 1 | V225G | Rv0402c | mmpL1 |
| Moldova | 32-12 | 565334 | 3003 | 83 | 1 | 0,0,43,40 | ATC | ACC | | 1 | I105T | Rv0474 | Rv0474 |
| Moldova | 32-12 | 3297146 | 424 | 65 | 0.246154 | 29,20,10,6 | TCC | TTC | | 1 | S232F | Rv2947c | pks15 |
| Moldova | 32-12 | 4074497 | 2918 | 81 | 1 | 0,0,49,32 | ATG | ATC | | 2 | M288I | Rv3635 | Rv3635 |
| Moldova | 33-11 | 1096567 | 2158 | 140 | 0.8 | 13,15,93,19 | . | . | . | | . | . | na |
| Moldova | 33-12 | 1096567 | 999 | 42 | 0.904762 | 2,2,31,7 | . | . | . | | . | . | na |
| Moldova | 33-12 | 3717105 | 2846 | 139 | 0.647482 | 19,30,36,54 | CAG | TAG | | 0 | Q6* | Rv3331 | sugI |
| Moldova | 34-11 | 2338275 | 4006 | 252 | 0.535714 | 92,25,106,29 | ACC | ACG | | 2 | T77T | Rv2081c | Rv2081c |
| Moldova | 34-12 | 2338275 | 1135 | 51 | 0.686275 | 13,3,23,12 | ACC | ACG | | 2 | T77T | Rv2081c | Rv2081c |
| Moldova | 35-11 | 4216118 | 2226 | 436 | 0.208716 | 145,200,39,52 | . | . | . | | . | . | na |
| Moldova | 35-12 | 4216118 | 2893 | 152 | 0.605263 | 22,38,39,53 | . | . | . | | . | . | na |
| Moldova | 39-13 | 2517465 | 525 | 65 | 0.307692 | 20,25,10,10 | GGG | AGG | | 0 | G227R | Rv2243 | fabD |
| Moldova | 39-13 | 2517868 | 308 | 82 | 0.170732 | 34,34,8,6 | TTC | TCC | | 1 | F33S | Rv2244 | acpM |
| Moldova | 40-11 | 454295 | 6852 | 258 | 0.94186 | 0,0,154,89 | CCA | CCG | | 2 | P26P | Rv0376c | Rv0376c |
| Moldova | 40-11 | 819912 | 910 | 315 | 0.136508 | 141,130,22,21 | GTG | GTA | | 2 | V196V | Rv0727c | fucA |
| Moldova | 40-11 | 1011865 | 306 | 338 | 0.047337 | 143,179,8,8 | TTC | TTT | | 2 | F45F | Rv0908 | ctpE |
| Moldova | 40-11 | 1254318 | 10550 | 385 | 0.825974 | 23,44,116,202 | GCG | ACG | | 0 | A73T | Rv1129c | Rv1129c |
| Moldova | 40-11 | 1834295 | 1439 | 369 | 0.173442 | 169,136,32,32 | GTC | TTC | | 0 | V252F | Rv1630 | rpsA |
| Moldova | 40-11 | 2289213 | 9367 | 346 | 0.815029 | 25,39,141,141 | CAG | CGG | | 1 | Q10R | Rv2043c | pncA |
| Moldova | 40-11 | 2318755 | 923 | 331 | 0.154079 | 112,168,19,32 | ATC | GTC | | 0 | I667V | Rv2062c | cobN |
| Moldova | 40-11 | 2815150 | 498 | 297 | 0.080808 | 121,152,12,12 | ATT | ATG | | 2 | I577M | Rv2501c | accA1 |
| Moldova | 40-11 | 3218303 | 7155 | 381 | 0.622047 | 73,69,123,114 | . | . | . | | . | . | na |
| Moldova | 40-11 | 3401002 | 6906 | 268 | 0.839552 | 18,24,100,125 | ACC | ACT | | 2 | T19T | Rv3040c | Rv3040c |
| Moldova | 40-11 | 3486940 | 369 | 368 | 0.057065 | 129,218,6,15 | GGA | GGG | | 2 | G144G | Rv3121 | cyp141 |

| | | | | | | | | | | | | | |
|---|---|---|---|---|---|---|---|---|---|---|---|---|---|
| Moldova | 40-12 | 454295 | 1533 | 55 | 0.981818 | 0,0,29,25 | CCA | CCG | | 2 | P26P | Rv0376c | Rv0376c |
| Moldova | 40-12 | 819912 | 417 | 96 | 0.177083 | 34,45,9,8 | GTG | GTA | | 2 | V196V | Rv0727c | fucA |
| Moldova | 40-12 | 1011865 | 5374 | 525 | 0.369524 | 147,184,80,114 | TTC | TTT | | 2 | F45F | Rv0908 | ctpE |
| Moldova | 40-12 | 1254318 | 12009 | 380 | 0.913158 | 13,20,157,190 | GCG | ACG | | 0 | A73T | Rv1129c | Rv1129c |
| Moldova | 40-12 | 1834295 | 573 | 246 | 0.109756 | 127,92,17,10 | GTC | TTC | | 0 | V252F | Rv1630 | rpsA |
| Moldova | 40-12 | 2289213 | 8570 | 274 | 0.908759 | 12,13,102,147 | CAG | CGG | | 1 | Q10R | Rv2043c | pncA |
| Moldova | 40-12 | 2815150 | 446 | 269 | 0.085502 | 106,140,8,15 | ATT | ATG | | 2 | I577M | Rv2501c | accA1 |
| Moldova | 40-12 | 3218303 | 720 | 89 | 0.337079 | 28,31,14,16 | . | . | . | | . | . | na |
| Moldova | 40-12 | 3401002 | 1782 | 52 | 0.980769 | 0,1,25,26 | ACC | ACT | | 2 | T19T | Rv3040c | Rv3040c |
| Moldova | 40-12 | 3486940 | 750 | 472 | 0.080508 | 203,231,15,23 | GGA | GGG | | 2 | G144G | Rv3121 | cyp141 |
| Moldova | 40-13 | 454295 | 1459 | 63 | 0.904762 | 0,0,33,24 | CCA | CCG | | 2 | P26P | Rv0376c | Rv0376c |
| Moldova | 40-13 | 819912 | 501 | 83 | 0.240964 | 34,29,10,10 | GTG | GTA | | 2 | V196V | Rv0727c | fucA |
| Moldova | 40-13 | 1011865 | 834 | 298 | 0.124161 | 107,154,15,22 | TTC | TTT | | 2 | F45F | Rv0908 | ctpE |
| Moldova | 40-13 | 1254318 | 6993 | 264 | 0.80303 | 24,28,87,125 | GCG | ACG | | 0 | A73T | Rv1129c | Rv1129c |
| Moldova | 40-13 | 1834295 | 715 | 125 | 0.24 | 54,41,14,16 | GTC | TTC | | 0 | V252F | Rv1630 | rpsA |
| Moldova | 40-13 | 2289213 | 5548 | 222 | 0.77027 | 22,28,78,93 | CAG | CGG | | 1 | Q10R | Rv2043c | pncA |
| Moldova | 40-13 | 2318755 | 832 | 85 | 0.376471 | 22,30,15,17 | ATC | GTC | | 0 | I667V | Rv2062c | cobN |
| Moldova | 40-13 | 2815150 | 295 | 235 | 0.068085 | 101,118,7,9 | ATT | ATG | | 2 | I577M | Rv2501c | accA1 |
| Moldova | 40-13 | 3218303 | 877 | 80 | 0.4 | 23,25,16,16 | . | . | . | | . | . | na |
| Moldova | 40-13 | 3401002 | 899 | 46 | 0.673913 | 7,8,17,14 | ACC | ACT | | 2 | T19T | Rv3040c | Rv3040c |
| Moldova | 40-13 | 3486940 | 592 | 330 | 0.09697 | 144,154,15,17 | GGA | GGG | | 2 | G144G | Rv3121 | cyp141 |
| Moldova | 40-14 | 454295 | 1021 | 44 | 0.909091 | 0,0,22,18 | CCA | CCG | | 2 | P26P | Rv0376c | Rv0376c |
| Moldova | 40-14 | 819912 | 218 | 56 | 0.160714 | 22,25,5,4 | GTG | GTA | | 2 | V196V | Rv0727c | fucA |
| Moldova | 40-14 | 1011865 | 3315 | 406 | 0.305419 | 110,172,47,77 | TTC | TTT | | 2 | F45F | Rv0908 | ctpE |
| Moldova | 40-14 | 1254318 | 9740 | 304 | 0.927632 | 7,15,122,160 | GCG | ACG | | 0 | A73T | Rv1129c | Rv1129c |
| Moldova | 40-14 | 1834295 | 208 | 201 | 0.054726 | 103,87,7,4 | GTC | TTC | | 0 | V252F | Rv1630 | rpsA |
| Moldova | 40-14 | 2289213 | 7048 | 237 | 0.869198 | 16,14,85,121 | CAG | CGG | | 1 | Q10R | Rv2043c | pncA |
| Moldova | 40-14 | 2815150 | 840 | 248 | 0.16129 | 78,130,15,25 | ATT | ATG | | 2 | I577M | Rv2501c | accA1 |
| Moldova | 40-14 | 3218303 | 861 | 77 | 0.402597 | 20,26,14,17 | . | . | . | | . | . | na |
| Moldova | 40-14 | 3401002 | 998 | 38 | 0.842105 | 3,3,12,20 | ACC | ACT | | 2 | T19T | Rv3040c | Rv3040c |
| Moldova | 40-14 | 3486940 | 1293 | 323 | 0.170279 | 132,136,22,33 | GGA | GGG | | 2 | G144G | Rv3121 | cyp141 |
| Moldova | 41-11 | 3596299 | 10270 | 332 | 0.921687 | 5,19,130,176 | GAG | GAA | | 2 | E412E | Rv3220c | Rv3220c |
| Moldova | 41-12 | 1133240 | 500 | 31 | 0.580645 | 7,5,9,9 | TCC | TTC | | 1 | S539F | Rv1013 | pks16 |
| Moldova | 43-13 | 1481185 | 10232 | 649 | 0.523883 | 1.56151E+11 | GAT | GAG | | 2 | D439E | Rv1319c | Rv1319c |
| Moldova | 45-11 | 760729 | 913 | 437 | 0.10984 | 195,193,26,22 | TAT | TGT | | 1 | Y308C | Rv0667 | rpoB |
| Moldova | 45-11 | 1326482 | 1282 | 541 | 0.107209 | 223,260,24,34 | GAC | GGC | | 1 | D344G | Rv1185c | fadD21 |
| Moldova | 45-11 | 3431783 | 1200 | 448 | 0.120536 | 183,211,24,30 | GTC | GCC | | 1 | V119A | Rv3067 | Rv3067 |
| Moldova | 46-11 | 1276785 | 16534 | 774 | 0.687339 | 1.09131E+11 | AGC | GGC | | 0 | S322G | Rv1148c | Rv1148c |
| Moldova | 46-11 | 2742077 | 3922 | 284 | 0.538732 | 64,58,78,75 | GGG | AGG | | 0 | G457R | Rv2443 | dctA |
| Moldova | 46-11 | 3836358 | 4557 | 325 | 0.483077 | 97,71,89,68 | TCC | TCT | | 2 | S178S | Rv3417c | groEL1 |
| Moldova | 46-12 | 46877 | 2637 | 173 | 0.508671 | 39,46,45,43 | GTG | CTG | | 0 | V111L | Rv0042c | Rv0042c |
| Moldova | 46-12 | 1276785 | 3181 | 186 | 0.580645 | 39,39,47,61 | AGC | GGC | | 0 | S322G | Rv1148c | Rv1148c |
| Moldova | 46-12 | 2186365 | 842 | 138 | 0.23913 | 61,44,20,13 | GGG | GGA | | 2 | G265G | Rv1935c | echA13 |
| Moldova | 46-14 | 1034862 | 745 | 156 | 0.211538 | 68,55,19,14 | . | . | . | | . | . | na |
| Moldova | 46-14 | 1276785 | 2799 | 157 | 0.624204 | 29,30,39,59 | AGC | GGC | | 0 | S322G | Rv1148c | Rv1148c |
| Moldova | 47-11 | 8208 | 10330 | 373 | 0.823056 | 31,35,139,168 | GAG | CAG | | 0 | E303Q | Rv0006 | gyrA |
| Moldova | 47-11 | 569897 | 1541 | 154 | 0.448052 | 29,53,32,37 | . | . | . | | . | . | na |
| Moldova | 47-11 | 1468208 | 6620 | 340 | 0.852941 | 37,13,128,162 | CTG | CGG | | 1 | L433R | Rv1313c | Rv1313c |
| Moldova | 47-12 | 8208 | 10944 | 304 | 0.996711 | 1,0,136,167 | GAG | CAG | | 0 | E303Q | Rv0006 | gyrA |
| Moldova | 47-12 | 569897 | 746 | 72 | 0.486111 | 15,21,20,15 | . | . | . | | . | . | na |
| Moldova | 47-12 | 1468208 | 3636 | 188 | 0.851064 | 20,8,62,98 | CTG | CGG | | 1 | L433R | Rv1313c | Rv1313c |
| Moldova | 48-11 | 1276588 | 777 | 380 | 0.121053 | 167,167,34,12 | GCG | GCC | | 2 | A387A | Rv1148c | Rv1148c |
| Moldova | 48-11 | 3235441 | 1061 | 194 | 0.216495 | 74,78,17,25 | GCC | GTC | | 1 | A789V | Rv2922c | smc |
| Moldova | 48-12 | 1276588 | 954 | 172 | 0.284884 | 63,60,38,11 | GCG | GCC | | 2 | A387A | Rv1148c | Rv1148c |
| Moldova | 48-13 | 1276588 | 1104 | 156 | 0.326923 | 55,50,36,15 | GCG | GCC | | 2 | A387A | Rv1148c | Rv1148c |
| Moldova | 48-14 | 50557 | 2074 | 66 | 0.924242 | 0,0,23,38 | AGG | GGG | | 0 | R190G | Rv0046c | ino1 |
| Moldova | 48-14 | 1276588 | 776 | 139 | 0.266187 | 52,50,29,8 | GCG | GCC | | 2 | A387A | Rv1148c | Rv1148c |
| Moldova | 51-11 | 1253819 | 1255 | 335 | 0.164179 | 135,145,32,23 | GCC | GTC | | 1 | A239V | Rv1129c | Rv1129c |
| Moldova | 51-11 | 1924420 | 481 | 403 | 0.062035 | 149,229,11,14 | CTC | GTC | | 0 | L198V | Rv1699 | pyrG |
| Moldova | 51-11 | 4088467 | 1200 | 308 | 0.178571 | 142,111,34,21 | GAC | GGC | | 1 | D22G | Rv3648c | cspA |
| Moldova | 51-12 | 1924420 | 17177 | 637 | 0.806907 | 61,62,242,272 | CTC | GTC | | 0 | L198V | Rv1699 | pyrG |

| | | | | | | | | | | | | |
|---|---|---|---|---|---|---|---|---|---|---|---|---|
| Moldova | 51-13 | 1924420 | 12434 | 452 | 0.818584 | 42,40,166,204 | CTC | GTC | 0 | L198V | Rv1699 | pyrG |
| Moldova | 52-11 | 412061 | 1611 | 331 | 0.193353 | 142,125,33,31 | GGC | GGG | 2 | G408G | Rv0342 | iniA |
| Moldova | 52-11 | 1253483 | 4356 | 322 | 0.459627 | 90,84,80,68 | GCG | GTG | 1 | A351V | Rv1129c | Rv1129c |
| Moldova | 52-11 | 1359880 | 3159 | 393 | 0.305344 | 159,114,72,48 | GCG | GCT | 2 | A89A | Rv1216c | Rv1216c |
| Moldova | 52-11 | 1692117 | 1725 | 335 | 0.226866 | 140,119,40,36 | CCT | CCG | 2 | P76P | Rv1501 | Rv1501 |
| Moldova | 52-11 | 1918406 | 3772 | 376 | 0.369681 | 103,134,63,76 | TCG | TGG | 1 | S156W | Rv1694 | tlyA |
| Moldova | 52-11 | 1918449 | 7321 | 375 | 0.637333 | 59,77,104,135 | TGC | TGA | 2 | C170* | Rv1694 | tlyA |
| Moldova | 52-12 | 717056 | 1615 | 46 | 0.978261 | 0,1,27,18 | CCC | CCT | 2 | P131P | Rv0624 | vapC30 |
| Moldova | 52-12 | 1918449 | 3954 | 113 | 0.982301 | 1,1,62,49 | TGC | TGA | 2 | C170* | Rv1694 | tlyA |
| Moldova | 52-12 | 2849510 | 2578 | 79 | 0.974684 | 1,1,39,38 | . | . | . | . | . | na |
| Moldova | 52-12 | 3422977 | 6335 | 190 | 0.952632 | 3,6,80,101 | TGC | TGT | 2 | C79C | Rv3060c | Rv3060c |
| Moldova | 53-11 | 2889376 | 2112 | 402 | 0.208955 | 163,155,43,41 | GCC | TCC | 0 | A1002S | Rv2566 | Rv2566 |
| Moldova | 53-12 | 2889376 | 655 | 119 | 0.210084 | 45,49,14,11 | GCC | TCC | 0 | A1002S | Rv2566 | Rv2566 |
| Moldova | 54-11 | 1096567 | 688 | 68 | 0.735294 | 8,10,42,8 | . | . | . | . | . | na |
| Moldova | 54-11 | 1161026 | 5070 | 171 | 0.953216 | 4,4,92,71 | GGC | GGT | 2 | G42G | Rv1038c | esxJ |
| Moldova | 54-12 | 1096567 | 632 | 41 | 0.804878 | 4,4,29,4 | . | . | . | . | . | na |
| Moldova | 54-12 | 1161026 | 3577 | 142 | 0.894366 | 7,8,74,53 | GGC | GGT | 2 | G42G | Rv1038c | esxJ |
| Moldova | 54-12 | 3015022 | 898 | 180 | 0.211111 | 69,73,18,20 | . | . | . | . | . | na |
| Moldova | 54-13 | 764328 | 3415 | 155 | 0.716129 | 23,21,47,64 | ATC | ACC | 1 | I320T | Rv0668 | rpoC |
| Moldova | 54-13 | 1096567 | 867 | 48 | 0.875 | 3,3,39,3 | . | . | . | . | . | na |
| Moldova | 54-13 | 1161026 | 3548 | 133 | 0.924812 | 5,5,71,52 | GGC | GGT | 2 | G42G | Rv1038c | esxJ |
| Moldova | 54-13 | 3861669 | 790 | 97 | 0.298969 | 26,42,11,18 | TCG | TCA | 2 | S94S | Rv3442c | rpsI |
| Moldova | 55-11 | 1340688 | 4900 | 689 | 0.323657 | 255,210,154,69 | CAC | CAT | 2 | H10H | Rv1197 | esxK |
| Moldova | 55-11 | 4393010 | 4369 | 436 | 0.355505 | 142,139,80,75 | GCT | CCT | 0 | A22P | Rv3907c | pcnA |

**Suplemnetary table 5**

**Position and frequency of the confident mixed SNPs identifed (in serial isolates) in genes with any reported link to antimycobactetial drug resistance in the literature**

| Site | Sample | Postion | Quality | ? | Proprtion mutant SNP | Base called frowar | Codon Wil | Codon mutant | | AA change | Gene | Gene name |
|---|---|---|---|---|---|---|---|---|---|---|---|---|
| Belarus | 01-06 | 777262 | 3583 | 302 | 0.430464 | 85,87,62,68 | ACC | CCC | 0 | T407P | Rv0676c | mmpL5 |
| Belarus | 02-06 | 4138521 | 2985 | 408 | 0.306373 | 170,111,71,54 | ATG | ACG | 1 | M412T | Rv3696c | glpK |
| Belarus | 08-06 | 495753 | 5392 | 264 | 0.666667 | 42,46,79,97 | GCC | GTC | 1 | A521V | Rv0410c | pknG |
| Belarus | 08-07 | 495753 | 5008 | 305 | 0.557377 | 68,67,78,92 | GCC | GTC | 1 | A521V | Rv0410c | pknG |
| Belarus | 08-07 | 276505 | 3685 | 369 | 0.371274 | 137,95,84,53 | CTT | TTT | 0 | L150F | Rv0231 | fadE4 |
| Belarus | 010-06 | 600096 | 7074 | 267 | 0.827715 | 22,24,113,108 | GCC | GCG | 2 | A966A | Rv0507 | mmpL2 |
| Belarus | 010-07 | 600096 | 8484 | 252 | 0.984127 | 2,2,127,121 | GCC | GCG | 2 | A966A | Rv0507 | mmpL2 |
| Belarus | 010-08 | 600096 | 731 | 269 | 0.130112 | 122,112,12,23 | GCC | GCG | 2 | A966A | Rv0507 | mmpL2 |
| Belarus | 010-06 | 3525768 | 8068 | 285 | 0.852632 | 21,21,130,113 | CGG | CTG | 1 | R546L | Rv3157 | nuoM |
| Belarus | 010-07 | 3525768 | 9335 | 264 | 0.988636 | 1,2,140,121 | CGG | CTG | 1 | R546L | Rv3157 | nuoM |
| Belarus | 010-08 | 3525768 | 811 | 323 | 0.114551 | 144,142,16,21 | CGG | CTG | 1 | R546L | Rv3157 | nuoM |
| Belarus | 11-08 | 761155 | 3326 | 374 | 0.336898 | 119,129,73,53 | TCG | TTG | 1 | S450L | Rv0667 | rpoB |
| Moldova | 1-12 | 3717108 | 649 | 106 | 0.254717 | 33,46,12,15 | CCT | GCT | 0 | P7A | Rv3331 | sugI |
| Moldova | 9-12 | 3717108 | 1843 | 100 | 0.59 | 16,25,22,37 | CCT | ACT | 0 | P7T | Rv3331 | sugI |
| Moldova | 11-13 | 8416 | 1763 | 316 | 0.224684 | 98,146,24,47 | GTC | GCC | 1 | V372A | Rv0006 | gyrA |
| Moldova | 16-11 | 778994 | 1149 | 119 | 0.10149 | 46,32,23,18 | AGC | ATC | 1 | S2I | Rv0678 | Rv0678 |
| Moldova | 16-11 | 2288920 | 10567 | 419 | 0.761337 | 43,57,148,171 | GGA | CGA | 0 | G108R | Rv2043c | pncA |
| Moldova | 16-11 | 2289081 | 468 | 412 | 0.058252 | 224,164,10,14 | CCG | CTG | 1 | P54L | Rv2043c | pncA |
| Moldova | 16-12 | 778994 | 745 | 335 | 0.344538 | 172,129,19,15 | AGC | ATC | 1 | S2I | Rv0678 | Rv0678 |
| Moldova | 16-12 | 2288920 | 2364 | 133 | 0.56391 | 33,25,37,38 | GGA | CGA | 0 | G108R | Rv2043c | pncA |
| Moldova | 16-12 | 2289081 | 1616 | 119 | 0.470588 | 34,29,31,25 | CCG | CTG | 1 | P54L | Rv2043c | pncA |
| Moldova | 22-12 | 596957 | 1382 | 208 | 0.264423 | 72,81,32,23 | GGC | AGC | 0 | G67S | Rv0506 | mmpS2 |
| Moldova | 26-12 | 2095631 | 3927 | 352 | 0.397727 | 101,111,70,70 | GTC | CTC | 0 | V180L | Rv1845c | blaR |
| Moldova | 32-11 | 482558 | 14110 | 455 | 0.896703 | 22,25,213,195 | GTC | GGC | 1 | V225G | Rv0402c | mmpL1 |
| Moldova | 32-12 | 482558 | 6954 | 197 | 0.994924 | 1,0,94,102 | GTC | GGC | 1 | V225G | Rv0402c | mmpL1 |
| Moldova | 33-12 | 3717105 | 2846 | 139 | 0.647482 | 19,30,36,54 | CAG | TAG | 0 | Q6* | Rv3331 | sugI |
| Moldova | 40-11 | 1834295 | 1439 | 369 | 0.173442 | 169,136,32,32 | GTC | TTC | 0 | V252F | Rv1630 | rpsA |
| Moldova | 40-11 | 2289213 | 9367 | 346 | 0.815029 | 25,39,141,141 | CAG | CGG | 1 | Q10R | Rv2043c | pncA |
| Moldova | 40-12 | 1834295 | 573 | 246 | 0.109756 | 127,92,17,10 | GTC | TTC | 0 | V252F | Rv1630 | rpsA |
| Moldova | 40-12 | 2289213 | 8570 | 274 | 0.908759 | 12,13,102,147 | CAG | CGG | 1 | Q10R | Rv2043c | pncA |
| Moldova | 40-13 | 1834295 | 715 | 125 | 0.24 | 54,41,14,16 | GTC | TTC | 0 | V252F | Rv1630 | rpsA |
| Moldova | 40-13 | 2289213 | 5548 | 222 | 0.77027 | 22,28,78,93 | CAG | CGG | 1 | Q10R | Rv2043c | pncA |
| Moldova | 40-14 | 1834295 | 208 | 201 | 0.054726 | 103,87,7,4 | GTC | TTC | 0 | V252F | Rv1630 | rpsA |

| | | | | | | | | | | | | |
|---|---|---|---|---|---|---|---|---|---|---|---|---|
| Moldova | 40-14 | 2289213 | 7048 | 237 | 0.869198 | 16,14,85,121 | CAG | CGG | 1 | Q10R | Rv2043c | pncA |
| Moldova | 45-11 | 760729 | 913 | 437 | 0.10984 | 195,193,26,22 | TAT | TGT | 1 | Y308C | Rv0667 | rpoB |
| Moldova | 45-11 | 1326482 | 1282 | 541 | 0.107209 | 223,260,24,34 | GAC | GGC | 1 | D344G | Rv1185c | fadD21 |
| Moldova | 47-11 | 8208 | 10330 | 373 | 0.823056 | 31,35,139,168 | GAG | CAG | 0 | E303Q | Rv0006 | gyrA |
| Moldova | 47-12 | 8208 | 10944 | 304 | 0.996711 | 1,0,136,167 | GAG | CAG | 0 | E303Q | Rv0006 | gyrA |
| Moldova | 51-11 | 1924420 | 481 | 403 | 0.062035 | 149,229,11,14 | CTC | GTC | 0 | L198V | Rv1699 | pyrG |
| Moldova | 51-12 | 1924420 | 17177 | 637 | 0.806907 | 61,62,242,272 | CTC | GTC | 0 | L198V | Rv1699 | pyrG |
| Moldova | 51-13 | 1924420 | 12434 | 452 | 0.818584 | 42,40,166,204 | CTC | GTC | 0 | L198V | Rv1699 | pyrG |
| Moldova | 52-11 | 1918406 | 3772 | 376 | 0.369681 | 103,134,63,76 | TCG | TGG | 1 | S156W | Rv1694 | tlyA |
| Moldova | 52-11 | 1918449 | 7321 | 375 | 0.637333 | 59,77,104,135 | TGC | TGA | 2 | C170* | Rv1694 | tlyA |
| Moldova | 52-12 | 1918449 | 3954 | 113 | 0.982301 | 1,1,62,49 | TGC | TGA | 2 | C170* | Rv1694 | tlyA |
| Moldova | 54-13 | 764328 | 3415 | 155 | 0.716129 | 23,21,47,64 | ATC | ACC | 1 | I320T | Rv0668 | rpoC |

**Supplementary table 6**
**Variation detected in Rv0678**

| | | |
|---|---|---|
| Moldova | 25-11 | 779133 ins C |
| Moldova | 25-12 | 779133 ins C |
| Moldova | 25-13 | 779133 ins C |
| Moldova | 25-14 | 779133 ins C |
| Moldova | 50-11 | 779133 ins C |
| Moldova | 50-12 | 779133 ins C |
| Moldova | 52-11 | some reads with 779117 T ins |
| Moldova | 52-12 | 779117 T ins |
| Moldova | 52-13 | 779187 G ins |
